# The Landscape Of Circular RNA Expression In The Human Brain

**DOI:** 10.1101/500991

**Authors:** Akira Gokoolparsadh, Firoz Anwar, Irina Voineagu

**Author notes:** Corresponding author: Assoc. Prof. Irina Voineagu, School of Biotechnology and Biomolecular Sciences, University of New South Wales, Kensington, Sydney NSW 2052 Australia.

## Abstract

Circular RNAs (circRNAs) are enriched in the mammalian brain and are upregulated in response to neuronal differentiation and depolarisation. These RNA molecules, formed by non-canonical back-splicing, have both regulatory and translational potential. Here, we carried out an extensive characterisation of circRNA expression in the human brain, in nearly two hundred human brain samples, from both healthy individuals and autism cases. We identify hundreds of novel circRNAs and demonstrate that circRNAs are not expressed stochastically, but rather as major isoforms. We characterise inter-individual variability of circRNA expression in the human brain and show that inter-individual variability is less pronounced than variability between cerebral cortex and cerebellum. We also find that circRNA expression is dynamic during cellular maturation in brain organoids, but remains largely stable across the adult lifespan. Finally, we identify a circRNA co-expression module upregulated in autism samples, thereby adding another layer of complexity to the transcriptome changes observed in autism brain. These data provide a comprehensive catalogue of circRNAs as well as a deeper insight into their expression in the human brain, and are available as a free resource in browsable format at: http://www.voineagulab.unsw.edu.au/circ_rna

## INTRODUCTION

Circular RNAs (circRNAs) are an emerging class of RNAs formed by the non-sequential back-splicing of pre-messenger RNAs [1]. CircRNAs are expressed in a tissue-specific manner and in some cases are more efficiently generated than their linear cognate mRNAs [2, 3]. Although circRNAs are considered to be primarily non-coding molecules, a subset of circRNAs can be translated [4]. Due to their circular nature, circRNAs lack a 5’ cap structure and a poly-A tail, which in turn renders them resistant to enzymatic degradation. Consequently, circRNAs are exceptionally stable.

Several studies have demonstrated that across a range of mammalian tissues, circRNA expression is most abundant in the brain, with an overall enrichment in the cerebellum [5–7]. Similarly, in *Drosophila*, circRNA expression is enriched in the nervous system compared to other tissues [8]. In addition to tissue specificity, circRNAs also show dynamic expression during neuronal differentiation and depolarisation [6, 7, 9, 10], and are highly concentrated in synaptosomes [6], indicating that these molecules likely play a functional role in neurons.

The extent to which circRNAs influence the expression levels of their parental genes is yet to be elucidated, but at least two mechanisms are known to be at play: they can function as miRNA sponges [11, 12], and can increase the transcriptional efficiency of their parental genes [13]. The former mechanism is important for pluripotency and differentiation [14], as well as astrocyte activation [15]. Remarkably, the first study to investigate circRNA loss-of-function *in vivo* revealed that interactions between circRNAs and miRNAs are important for normal brain function [12]. *Cdr1as* knock-out mice, which lack circRNA expression at the *Cdr1as* locus, display impaired sensory-motor gating and abnormal synaptic transmission. These phenotypes are associated with changes in miR-7 and miR-671 levels, and consequent transcriptional changes in immediate early gene expression.

Despite accumulating evidence for an important role of circRNAs in the brain, our current understanding of their expression in the human brain is limited by relatively low sample sizes of existing studies. The largest study to date included 12 ENCODE normal human brain samples, while several other studies have assessed circRNA expression in small cohorts with at most 10 samples/group [5, 16–20]. A recent study combined data across multiple datasets, including a total of 21 human brain samples [21]; however, 19 of these samples were poly-A selected and thus circRNA detection was dependent on the inefficiency of the poly-A selection step. Therefore, a robust large-scale dataset of circRNA expression in the human brain is currently lacking.

Given the limited data on circRNA expression in the human brain, several questions remain open: (a) How does circRNA expression vary across individuals and human brain regions? (b) Does circRNA expression change significantly in an age-dependent manner? (c) Given that the brain is characterised by extensive alternative splicing (AS), is the abundant circRNA expression observed in the brain primarily a reflection of the complexity of brain AS events?

Here, we begin to address these questions by investigating circRNA expression in nearly two hundred human brain samples, from three brain regions: frontal cortex, temporal cortex and cerebellum, from normal individuals and autism spectrum disorder (ASD) cases. We first assessed the global properties of circRNA expression in control samples, to gain novel insights into the expression of this class of RNA molecules in the human brain.

- We identify hundreds of novel circRNAs, some of which are expressed in over a hundred brain samples.
- We demonstrate for the first time that circRNA expression is characterized by major isoforms, rather than stochastic expression.
- By investigating the interplay between AS and circRNA expression, we demonstrate that circRNAs are formed primarily from exons with high inclusion rates, supporting the notion that they are not primarily a by-product of exon skipping in the brain.
- We find that circRNAs only mildly change in expression in the human brain across the adult life span.
- We also investigate the developmental aspect of circRNA expression by assessing for the first time circRNA expression in human brain organoids [22], and providing the first resource of circRNA expression in brain organoids.

CircRNA expression has been previously shown to be brain-region specific for a subset of circRNAs [6]. Here, due to the dataset sample size we are able for the first time to investigate the relationship between inter-individual variability and brain-region specific circRNA expression. We show that similarly to protein-coding gene expression, inter-individual variability of circRNA expression is less pronounced than variability between brain regions (cerebral cortex and cerebellum), a property that holds true for control as well as ASD samples.

By investigating circRNA expression in ASD brain for the first time, we provide another layer of complexity to our understanding of transcriptome changes in ASD. Using a co-expression network approach, we identify circRNAs with increased expression in cerebral cortex in ASD.

Overall, our data provides a rich resource of circRNA expression in the human brain, and brings further insight into the expression properties of this class of RNA molecules.

## RESULTS

### Dataset Overview and Benchmarking

We investigated circRNA expression in a total of 197 human brain samples [23] from frontal cortex, temporal cortex and cerebellum (Methods; Supplementary Table 1) obtained from control (CTL) and ASD individuals (Figure 1a). These paired-end unstranded RNA-seq data were generated following ribosomal RNA-depletion (Methods; [23]), which makes them suitable for circRNA detection, unlike other large-scale brain datasets, which include poly-A selection (e.g. GTEX data [24]). In order to assess the reproducibility of our findings, we divided these samples into a discovery dataset (DS1, 144 samples) and a replication dataset (DS2, 53 samples). To allow adequate statistical power, the discovery dataset DS1 was assigned a larger sample size than the replication dataset DS2, where we would test for effects already identified in DS1. Given the well documented transcriptional similarity between frontal and temporal cortex [25], which we also observed for circRNAs (see results section on inter-region variability), we considered these regions as a single group (cerebral cortex; CTX). Within each dataset, there was no significant difference in age or gender ratios between CTX and cerebellum (CB) samples, or between ASD and CTL samples (Supplementary Figure 1).

**Figure 1.**
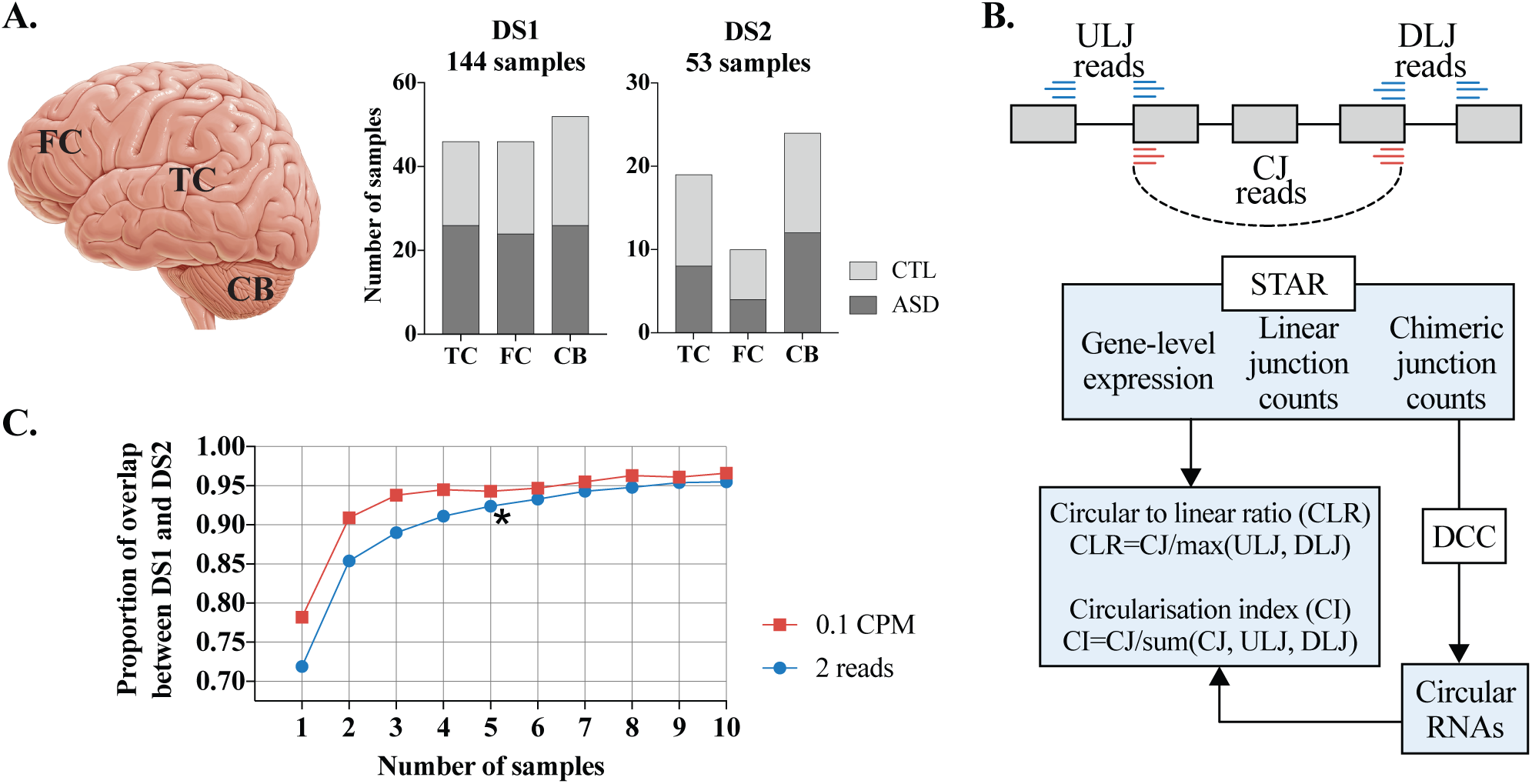
Dataset Overview. **(A)** Sample composition of dataset 1 (DS1) and dataset 2 (DS2). *Left:* schematic representation of brain regions included in the study. FC - frontal cortex; TC - temporal cortex; CB - cerebellum. *Right:* barplot displaying the number of samples included in DS1 and DS2 categorised by brain region and phenotype (ASD and CTL). **(B)** Outline of data analysis approach. *Top:* schematic representation of a gene expressing circRNA. Grey boxes - exons; grey lines - introns; blue lines - linear junction reads; red lines - circular (i.e. backsplice) junction reads. ULJ - upstream linear junction; DLJ - downstream linear junction. CJ - circular junction. *Bottom:* data analysis pipeline. **(C)** Overlap between circRNAs detected in DS1 and DS2 as a function of filtering parameters. y-axis: Overlap between DS1 and DS2 as a proportion of the smaller dataset (DS2); x-axis: number of independent samples in which detection (by either 2 back-splice junction reads or 0.1 CPM) is required. Based on these data, the chosen filtering parameter was 2 back-splice junction reads detected in a minimum of 5 independent samples (*).

To assess the quality of the RNA-seq data and of the circRNA quantification approach, we first carried out a benchmarking analysis:

- For a subset of 5 brain tissue samples we generated two sets of benchmarking data: poly-A+ RNA-seq (to assess false-positive rates of circRNA detection), and ribo-depleted stranded RNA-seq (to asses the effect of strandedness on circRNA quantification). These benchmarking data were generated from the same individual and brain region as the original data [23].
- We quantified circRNA expression using two different algorithms: CIRCexplorer [26] and DCC [27]. Based on a recent independent comparison of the performance of existing circRNA-quantification methods [28], both CIRCexplorer and DCC had a low false-positive rate, were robust to background noise, and had good sensitivity of detecting true positives (i.e. RNaseR enriched circRNAs).

The false-positive detection rate (i.e. the percentage of circRNAs detected in DS1 data that were also detected in the polyA+ data from the same sample, normalized for sequencing depth) was < 1% for DCC and < 3% for CIRCexplorer (Methods; Supplementary Figure 2a). Notably, we observed low false-positive rates despite the fact that the sequencing depth and the paired-end read length were higher for the polyA+ libraries than the original RNA-seq data (Supplementary Figure 2b). Since previously reported polyA+ based false positive rates were 2.7%-8% depending on algorithm [10], we concluded that both DCC and CIRCexplorer performed well on the brain dataset, with DCC showing a particularly low false-positive rate.

**Figure 2.**
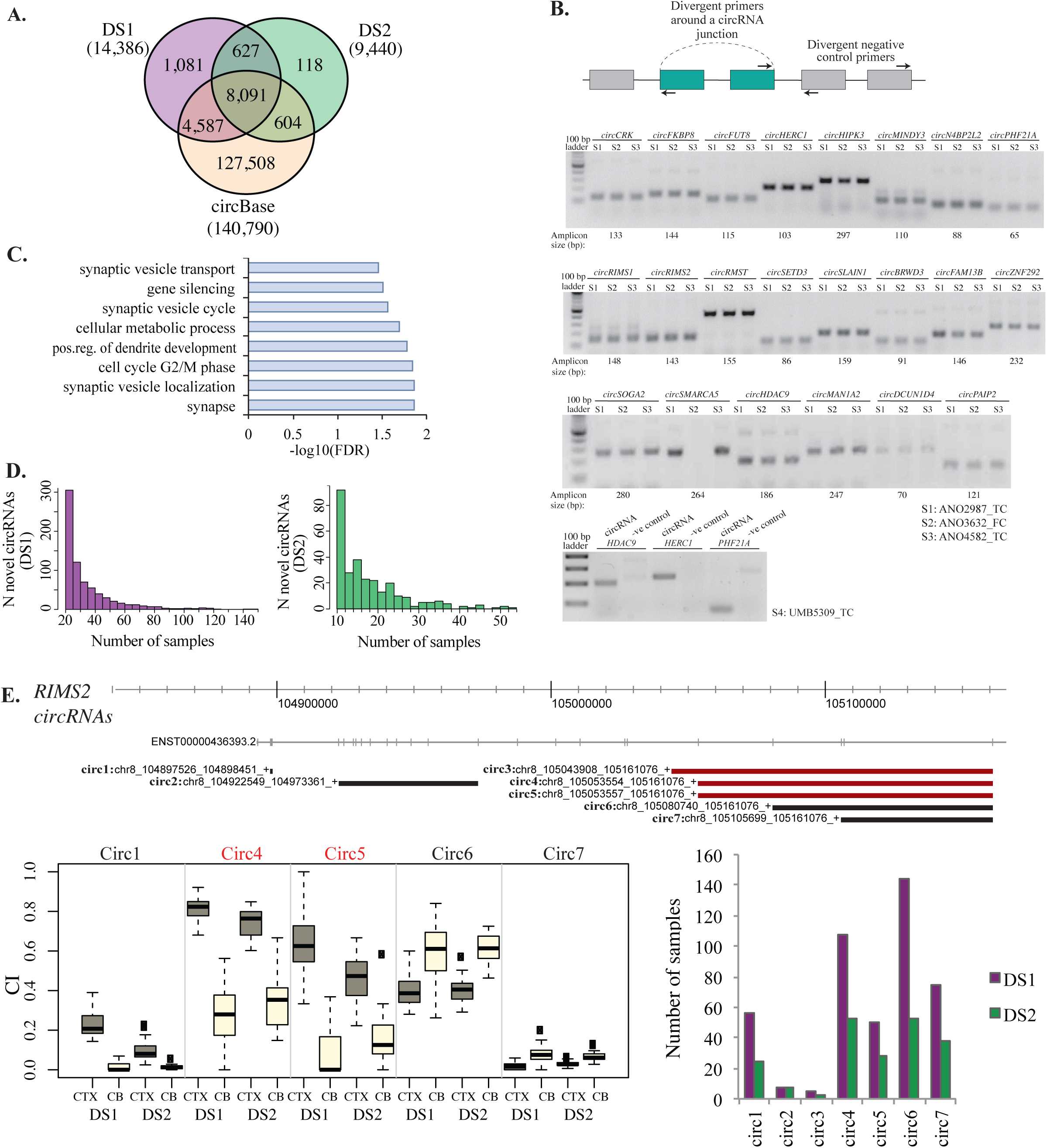
Dataset characterisation. **(A)** Venn diagram displaying the overlap between DS1, DS2 and circBase. The numbers between brackets show the total number of circRNAs in each dataset, after filtering for circRNAs expressed in a minimum of 5 samples in each dataset. **(B)** circRNA PCR validation Top: schematic display of divergent primers (black arrows) around a circRNA junction (dashed arc) which would amplify circRNAs, but not linear RNA molecules; negative control divergent primers anneal to exons that do not circularise, within the same gene. Bottom: Agarose gel electrophoresis of RT-PCR products using divergent primers around 22 selected circRNA junctions. Each RT-PCR is carried out on 3 brain samples (S1-S3). All circRNAs generate a PCR product at the expected size, except *circRMST*, and *circDCUN1D4*, for which the product is not of the expected size. Additional bands at higher length are likely rolling circle amplification products. For a subset of circRNAs, RT-PCR was carried out with primers around the circRNA junction, and negative control primers, on a distinct sample (S4; bottom panel). **(C)** Gene ontology enrichment of circRNA producing genes. FDR - false discovery rate. The most biologically relevant enriched terms are displayed. A full list of enriched terms is provided in Supplementary Table 3. **(D)** Histogram displaying novel circRNAs expressed in more than 10 samples. Left: DS1. Right: DS2. **(E)** Top: schematic display of seven highly expressed *RIMS2* circRNAs. Top track: hg19 chromosomal coordinates and a representative Ensembl transcript annotation of the region (due to space limit only one of multiple RIMS2 transcripts annotated at this position is displayed). Each circRNA is displayed as a line spanning the interval between its start and end junctions. Red: Novel circRNAs. Black: circRNAs present in circBase. Bottom, left: Boxplots displaying circRNA expression differences between CTX and CB, in both DS1 and DS2. CI: circularisation index. Only five of the seven circRNAs are displayed, which showed significant differences in CI levels between CTX and CB after correction for covariates and multiple testing (Supplementary Table 7). Boxplots were generated using the boxplot function in R; the horizontal line represents the median, boxes extend between the first and third quartiles, and whiskers extend from the box to 1.5x the inter-quartile range. Bottom, right: Barplot showing the number of samples in which each circRNA was detected. circRNA labels correspond to the labels from the top annotations track.

We also assessed the effect of strand-specificity of the RNA-seq data on false-positive rate detection. We found that all false-positive circRNAs detected in the unstranded data were detected in the stranded data as well (Supplementary Figure 2c), demonstrating that lack of strand-specificity did not lead to false-positive circRNA detection.

Across the five DS1 brain tissue samples, DCC and CIRCexplorer identified 5706 and 5426 circRNAs, respectively (Methods), with 91% of these being identified by both algorithms. Furthermore, the correlation between circRNA expression quantified by DCC and CIRCexplorer in the same sample was between 0.97 and 0.99 (Supplementary Figure 2d). This result indicated that circRNA quantification was robust to the choice of method, and due to its lower false-positive detection rate, DCC was chosen for downstream analyses.

CircRNA quantification in all 197 samples (including control and ASD samples) lead to the detection of a total of 43,872 circRNAs in DS1 and 28,251 circRNAs in DS2 (Methods). Circular junction (i.e. back-splice) reads were normalised to total library size as counts per million (CPM). Given that the expression of circRNAs can be influenced by the expression level of the parental transcript, we also normalised circRNA expression to that of the parental transcript by calculating a circular-to-linear ratio (CLR), as well as a circularisation index (CI; Figure 1b).

CircRNAs were considered robustly expressed if they were detected by at least 2 circular junction reads per sample in a minimum of 5 distinct samples. We selected these filtering criteria based on assessment of reproducibility between DS1 and DS2 at a range of filtering criteria (Methods, Figure 1c and Supplementary Figure 2e). Since our purpose was to construct a resource of circRNAs reliably detectable in the human brain, we considered reproducibility between independent sets of samples to be an important criterion. Requiring circRNA detection by at least 2 circular junction reads per sample in a minimum of 5 distinct samples maximised the reproducibility between DS1 and DS2 (90% overlap between datasets; Figure 2a).

The filtered datasets (DS1: 14,386 circRNAs, DS2: 9,440 circRNAs) were used for all downstream analyses.

We assessed the circular nature of a subset of 22 circRNAs by RT-PCR with divergent primers (which would amplify on a circular but not a linear molecule), and found a 90.9% validation rate (Methods; Figure 2b).

We also observed a strong correlation (Spearman *rho=0.93)* between the mean circRNA expression level (CLR) in DS1 and DS2, supporting the robustness of these data.

The 14,386 DS1 circRNAs and 9,440 DS2 circRNAs (listed in Supplementary Table 2) were expressed from a total of 4,555 and 3,650 unique genes, respectively. Gene ontology enrichment analysis of circRNA producing genes, with correction for gene length (Methods), showed an over-representation of genes functioning at the synapse, particularly those involved in synapse vesicle transport and localisation, but also genes involved in cell cycle G2/M phase transition, chromatin organisation and gene silencing (Figure 2c and Supplementary Table 3).

Previous data has shown that around 6% of the circRNAs detected in the human brain are also detected in mouse brain, with 28% of mouse circRNAs showing conserved expression in human [6]. We assessed the conservation rate of circRNA expression between human and mouse by using the liftOver tool [29] to obtain orthologous mouse coordinates of our circRNAs. The proportion of circRNAs for which the strict orthologous mouse coordinates were also detected as circRNAs was then assessed using mouse brain circRNA expression data from Rybak-Wolf et al. 2015 [6] (Methods). We found a circRNA expression conservation of 8.6% in DS1 and 9.6% in DS2, consistent with previous observations [6].

Overall the circRNA data showed high reproducibility across the two datasets, high RT-PCR validation rates, and were consistent with previous data regarding human-to-mouse conservation and expected functional enrichment of circRNA-producing genes.

### Identification of novel circRNAs

Given the much larger dataset employed in our study compared to existing human circRNA expression studies [6], which are curated in circBase [30], we expected to identify novel circRNAs despite our more stringent detection criteria (the higher stringency in our dataset comes from the fact we required detection of circRNAs in a minimum of 5 independent samples in each dataset, which had not been feasible with previous human brain sample sizes). Indeed, relative to circBase, we detected 1,548 novel circRNAs in DS1 and 692 in DS2, of which 83% were detected in both datasets (Methods; Figure 2a). Hundreds of novel circRNAs were detected in more than 10 brain samples, and some were expressed in more than a hundred samples, demonstrating that they are frequently expressed in the human brain (Figure 2d). CircRNA annotation relative to genomic features showed that most circRNAs, including novel circRNAs, were formed between annotated exon-exon junctions (Supplementary Figure 3).

**Figure 3.**
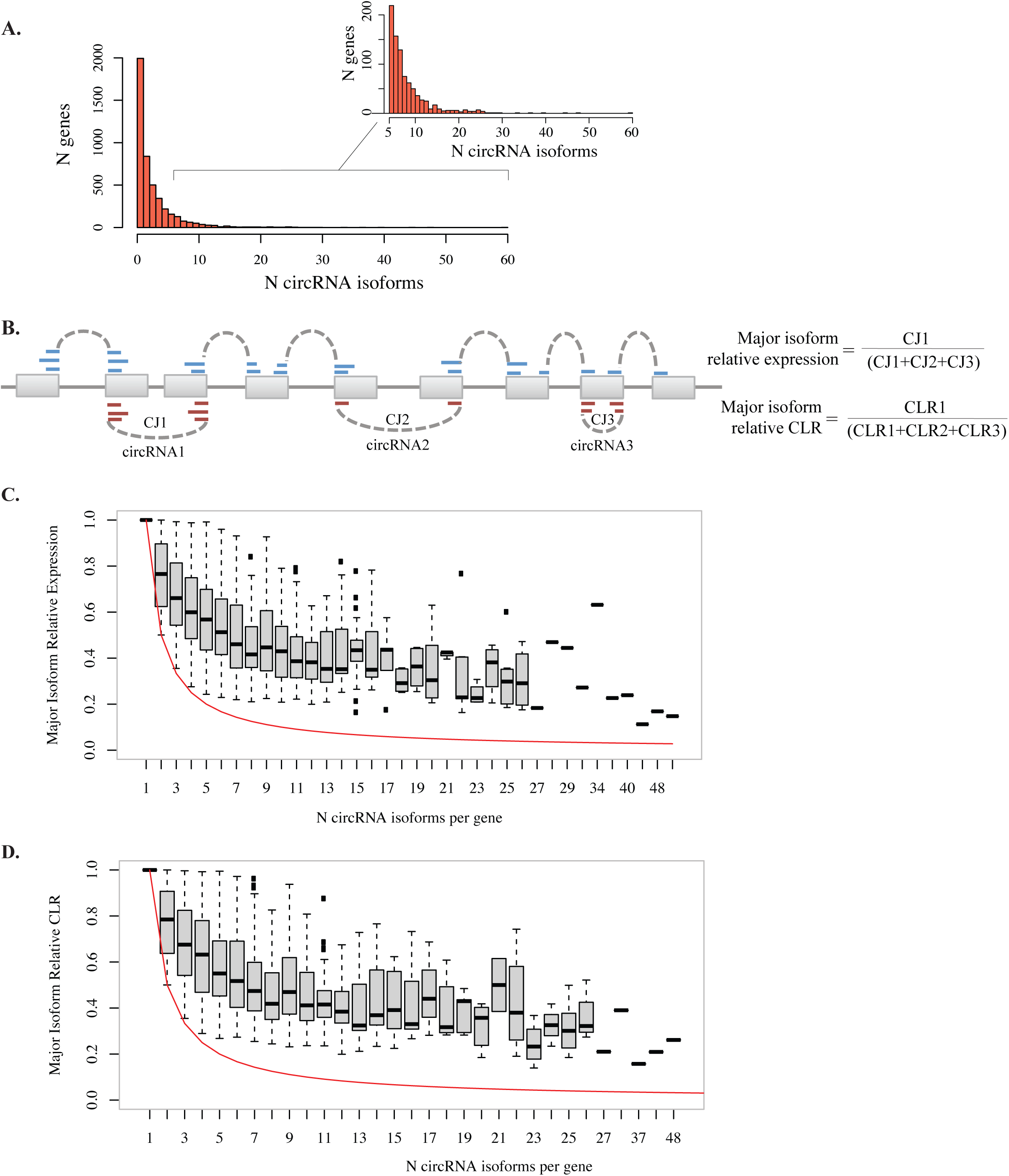
Major circRNA isoform expression. **(A)** Histogram of the number of circRNA isoforms per gene (DS1). *Left:* all genes. *Right:* genes with more than 5 isoforms per gene (blow-out of the data in the left panel). **(B)** *Top*: schematic representation of a hypothetical gene expressing three circRNAs (circRNA1–3), of which circRNA1 has the highest expression level. Circular junction (CJ) reads are shown in red; linear junction reads are shown in blue. Circular-to linear ratios (CLR) are calculated for each circRNA as shown in Figure 1. *Bottom:* Formulas for the major isoform relative expression and major isoform relative CLR for the hypothetical example shown in the top schematic. **(C)** and **(D)** Boxplots showing the major isoform relative expression and major isoform relative CLR respectively. Red line displays the expected major isoform relative expression and CLR if all isoforms were equally expressed. Boxplots were generated using the boxplot function in R; the horizontal line represents the median, boxes extend between the first and third quartiles, and whiskers extend from the box to 1.5x the inter-quartile range.

The novel circRNAs identified included both novel isoforms from known circRNA-producing genes, as well as over a thousand circRNAs expressed from genes not previously reported to circularise (Supplementary Table 2). Throughout this manuscript, we use the term “circRNA isoforms” to refer to circRNAs produced by back-splicing of distinct exon-exon junctions of a gene.

Among the novel circRNA-producing genes, several are involved in psychiatric disorders including *MEF2C-AS1* (a susceptibility locus for Alzheimer’s disease [31]) and *BRINP2* (a locus strongly associated with schizophrenia [32, 33]). Our data adds another layer to the transcriptome complexity of these genes, with potential implications for their transcriptional regulation.

As an example of the insight into circRNA expression in the human brain provided by our study, we outline circRNA expression from *RIMS2*, a gene that encodes a presynaptic protein involved in regulating synaptic membrane exocytosis [34]. *RIMS2* shows conserved circRNA expression in human, mouse, and pig brain [6, 35], with 35 known isoforms of *circRIMS2* in human brain [30]. Here, we identify 15 *RIMS2* circRNA isoforms detected in the human brain in both DS1 and DS2, of which 7 are novel circRNAs (Figure 2e). Notably, 3 of the novel circRNAs are highly expressed in both datasets (> 0.1 CPM in at least 2 distinct samples), and 2 of the novel isoforms were expressed in more than 50 brain samples. In addition to the high complexity of *circRIMS2* isoform expression, we also find that *RIMS2* shows circRNA isoform switching between cerebral cortex and cerebellum (Figure 2e).

### CircRNA expression is characterised by major isoform(s)

To investigate the global properties of circRNA expression in the human brain, we first used the data from control samples (N=68 in DS1; N=29 in DS2). Previous studies have shown that circRNA expression generally does not follow the expression of the parental gene [2, 6, 36]. Consistently, we found that circRNA expression levels did not correlate with that of the parental gene, whether the latter was measured as total gene-level expression, or as the expression of the corresponding linear junction (Spearman *rho:* 0.09 and 0.09 respectively for DS1; 0.08 and 0.06 for DS2; Supplementary Figure 4). This observation is generally interpreted to suggest that circRNA expression is regulated independently of their parental linear transcript expression. However, given that circRNAs are stable molecules, and thus are less susceptible to degradation than mRNAs [37, 38], the correlation between circRNA and mRNA might be attenuated due to the different degradation rates.

**Figure 4.**
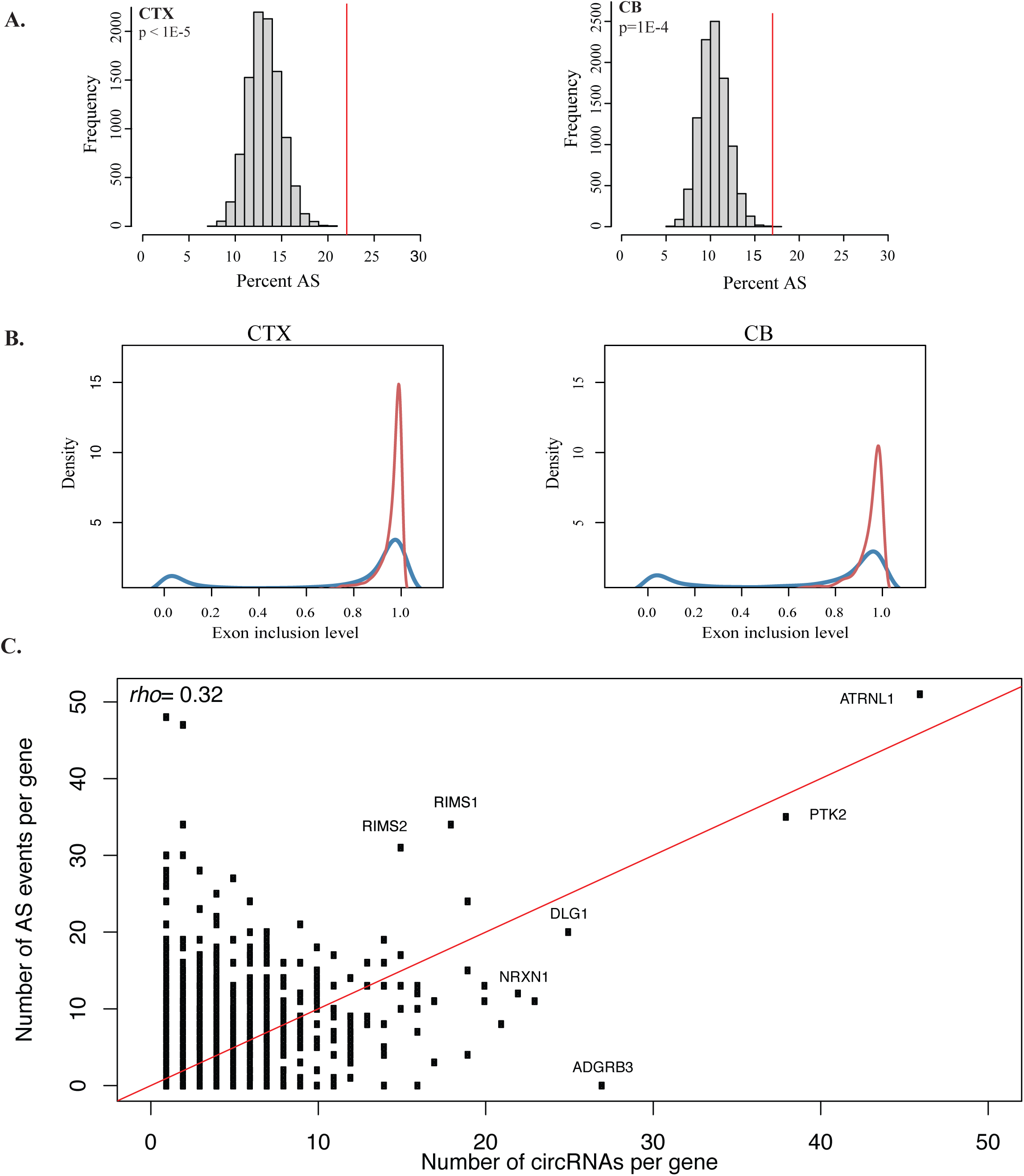
Interplay between circRNA expression and alternative splicing. **(A)** The percent of alternatively spliced exons (Percent AS) is significantly higher among circ-forming exons (red line) than among non-circ forming exons (histogram displaying percent AS for 10,000 random samplings of non-circ forming exons with similar flanking intron length as circ-forming exons, sampled from genes with similar expression levels as circ-forming genes). P-values were calculated as the number of random sampling scores with more extreme values than the circ-exons score. Left: CTX, DS1; right: CB, DS1. **(B)** Density plots of exon inclusion level for circ-exons (red) and non-circ exons (blue) in CTX (left) and CB (right). **(C)** Scatterplot of the number of alternative splicing (AS) events per gene (y-axis) and the number of circRNAs expressed per gene (x-axis). Each data point represents a gene. Gene symbols are displayed for the genes discussed in text.

To further investigate whether circRNAs show evidence of regulated expression in the human brain, we assessed the relative expression of circRNA isoforms for each circRNA-producing gene.

An important layer of regulation of mRNA expression consists of major isoform(s), which account for most of the transcriptional output of a gene in a given tissue of cell type. We thus asked whether circRNA isoforms are stochastically expressed, or show evidence of major isoform expression. The latter scenario would strongly indicate regulated circRNA expression, in a manner that is not confounded by the difference in circRNA vs. mRNA degradation rates (Figure 3).

We thus assessed the major circRNA isoform relative expression (i.e. the expression level of the most highly expressed circRNA relative to the total circRNA expression from a given gene; Figure 3b). We classified genes by the number of circRNAs expressed, and found that across a wide range of such classes, the major circRNA isoform accounts for the majority of circRNA output (Figure 3c). To test whether this simply reflects the linear major isoform expression, we carried out the same analysis using circular-to-linear ratios. Based on CLR data, we also found that the major circRNA isoform accounts for the majority of circRNA expression normalised to linear transcript expression (Figure 3d).

We then investigated whether major isoform expression is explained by sequence complementarity of flanking introns, i.e. does the major isoform tend to show the highest sequence complementarity? Reverse-complementary sequence matches (RCM) of flanking introns were calculated for all circRNAs using autoBLAST ([39]; Methods). We found that circRNA major isoforms were only marginally more likely than expected by chance to have the highest sequence complementarity score. For example, among genes producing 2 circRNAs, the major isoform showed the highest RCM score in only 53% of cases, while for genes producing 3 circRNAs the major isoform showed the highest RCM score in 36% of cases (Supplementary Figure 5).

**Figure 5.**
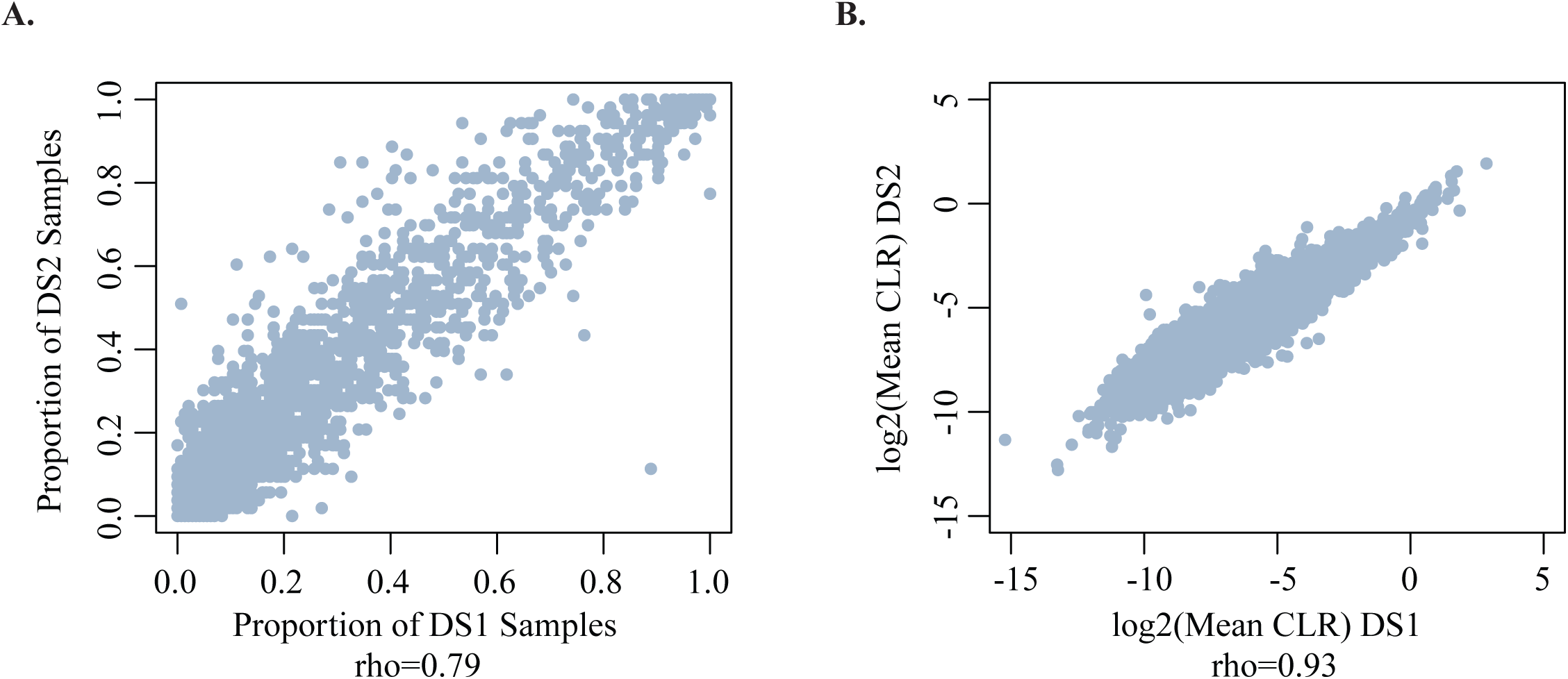
CircRNA expression variability. **(A)** Scatterplot showing the proportion of samples expressing a given circRNA at >= 0.1CPM in DS1 (x-axis) and DS2 (y-axis). Each data point represents a circRNA. Only circRNAs detected in both datasets are included. rho: Spearman correlation coefficient. **(B)** Scatterplot showing the mean CLR in DS1 (x-axis) and DS2 (y-axis). Each data point represents a circRNA. Only circRNAs detected in both datasets are included. rho: Spearman correlation coefficient.

These data demonstrate for the first time that circRNAs are expressed predominantly as major isoforms and further support the notion that circRNA formation is a regulated process.

### Interplay between canonical- and back-splicing in the human brain

To assess the interplay between alternative splicing (AS) and exon circularisation in the normal human brain, we characterised alternative splicing events in the larger dataset (DS1, control samples) using rMATS [40], and contrasting CTX and CB (Methods). Cassette exon (i.e. exon skipping) was the predominant AS event (72%), and thus we focused on the comparison of alternatively spliced cassette exons (referred to as “AS” from here on) and circRNA formation.

We found that the percentage of AS exons was significantly higher among circRNA-forming exons than among non-circ forming exons after correction for gene expression levels and intron length (Figure 4a).

To further investigate the relationship between alternative splicing and circularisation, we quantified exon inclusion level (i.e. the ratio between the inclusion and the skipping isoform of a given AS exon; Methods). When comparing exon inclusion levels between circ-exons and non-circ-exons that undergo alternative splicing, we found that circ-exons were formed primarily from exons with high inclusion rates (Figure 4b), indicating that circRNA formation is not primarily a by-product of exon skipping.

The number of circRNAs expressed per gene varied between one and sixty (DS1), and was highly correlated between DS1 and DS2 (Spearman *rho* = 0.82). Using circRNAs expressed in both DS1 and DS2, we found that 446 genes expressed more than 5 circRNAs. Given the observed association between circRNA expression and AS, we investigated whether these “circRNA hotspot genes” were also characterised by complex AS. We found a significant correlation between the number of circRNAs expressed and the number of alternative splicing events detected per gene *(rho=0.32*, p< 2.2e-16). The correlation remained significant after correction for the total number of exons per gene *(rho=0.14*, p< 2.2e-16). Some of the major hotspot genes (Figure 4c), including *RIMS1* (a schizophrenia associated gene [33]) and *NRXN1* (involved in autism [41, 42]) were indeed characterised by extensive alternative splicing (Figure 4c). However, hotspots of circRNA expression were also observed for 22 genes showing no detectable AS event (Figure 4c).

Overall, our data suggests that AS often co-occurs with circRNA formation, yet the interplay between AS and circRNA formation is highly locus-specific.

### Inter-individual variability of circRNA expression is less pronounced than inter-region variability

Transcriptome data from human brain tissue commonly shows that inter-individual variability of gene expression is less pronounced than the similarity within broad brain regions, such as cerebral cortex and cerebellum [25]. As a consequence, gene expression data commonly clusters by broad brain region, while cerebral cortex sub-regions, such as frontal and temporal cortex, are transcriptionally very similar and cluster together [25, 43].

To begin to understand the inter-individual variability of circRNA expression in the human brain, we first compared the relationship between mean expression and variance for canonical splice junctions and circRNAs (i.e. back-splice junctions) using the DS1 control samples (N=68). We observed a higher coefficient of variance [CV] for circRNA back-splice junctions compared to canonical splice junctions (mean CV: 3.6 and 3.08 respectively; p < 2.2e-16, Wilcoxon rank sum test; Supplementary Figure 6a). This difference is explained by the fact that while most splice junctions are detected in nearly all samples, circRNAs are often expressed in a small proportion of samples (Supplementary Figure 6b). This observation is consistent with (a) the fact that the efficiency of canonical splicing is higher than that of back-splicing, and consequently back-splicing occurs less frequently [9] and (b) the well documented property of non-coding RNAs to be specifically rather than broadly expressed [44].

**Figure 6.**
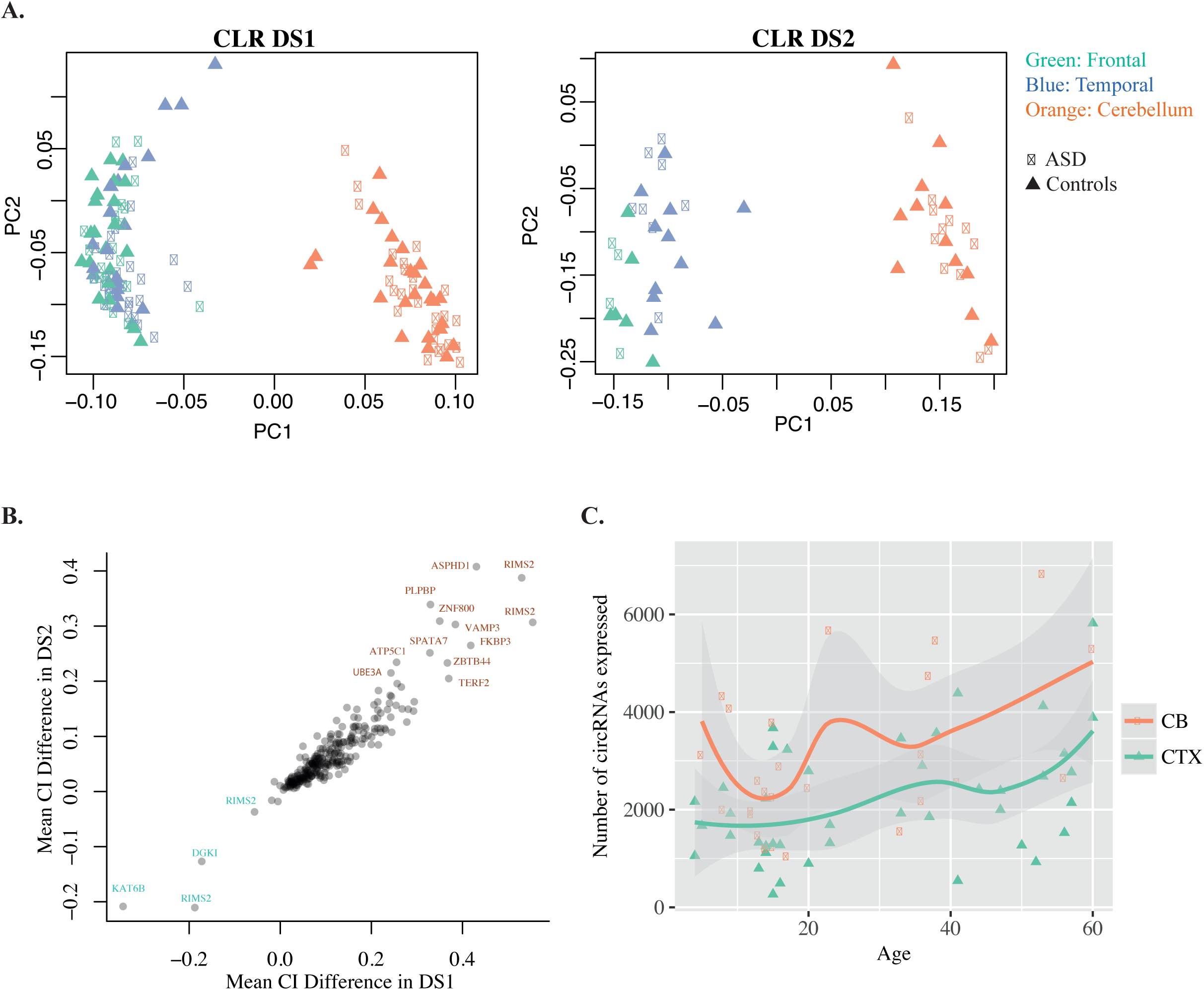
circRNA expression in CTX and CB. **(A)** Principal component plots of normalised circRNA expression (CLR) data. *Left:* DS1. *Right:* DS2. PC1, PC2 - first and second principal components. **(B)** Scatterplot of the circRNA expression difference between CB and CTX in DS1 (x-axis) and DS2 (y-axis). **(C)** Scatterplot displaying the total number of circRNAs expressed in each sample (y-axis; Methods) versus age (x-axis) in CB and CTX. Each data point represents a sample. Regression lines were generated using a loess smoothing function (geom_smooth) in ggplot2.

Since circRNA formation is a more rare event than the transcription of the parental transcript, we then investigated whether its frequency was consistent across the two datasets. We found high agreement between the proportion of samples in which a circRNA was detected in DS1 and DS2, as well as high correlation between CLRs in the two datasets (Figure 5a-b). These data indicate that although circRNA formation is rare, the frequency and expression level are intrinsic properties of circRNAs.

We also found that circRNA expression data clustered by brain region, even after normalisation of circRNA levels to the expression of the parental transcript (i.e. CLR), suggesting that similarly to mRNA expression, circRNA expression variability between individuals is less pronounced than region-specific differences (CTX and CB, Figure 6a). Within CTX, frontal and temporal cortex samples clustered together, as commonly observed for gene expression data. ASD samples did not show distinct clustering based on CLR, rather followed the overall pattern of clustering based on brain CTX and CB region (Figure 6a).

### CircRNA expression differences between CTX and CB

One of the important properties of human brain regions is their cellular composition and layer structure, which in turn can affect the cellular composition of dissected brain samples. Therefore, we estimated *in-silico* the proportion of individual cell types in the brain tissue samples, using DeconRNAseq [45] (Methods), and gene expression data from pure populations of immunopanned neurons, astrocytes, oligodendrocytes, microglia and endothelial cells [46] as reference transcriptomes. We found a significant difference in the proportion of neurons between CB and CTX (Supplementary Figure 7a), with lower and more homogeneous neuronal proportions in the CB samples. We also found that the first principle components of both gene expression and circRNA expression data strongly correlated with the estimated proportion of neurons (|*rho*| > 0.8 for gene expression PC1; *|rho|* > 0.6 for circRNA expression PC1; Supplementary Figure 7b-c), indicating that cellular composition is an important covariate when assessing gene and circRNA expression differences between brain regions. Thus we assessed circRNA expression differences between control CTX and CB samples using a linear model that included the estimated neuronal proportions as a covariate, in addition to age, sex, brain bank, RNA integrity number (RIN), and sequencing batch (Methods). We identified 501 circRNAs for which expression normalised to the linear transcript (i.e. CI) was significantly different between CTX and CB (FDR < 0.05; Methods). 266 of the 501 circRNAs were replicated as significant in DS2 (FDR < 0.05; Methods), with high agreement of directionality (Figure 6b). Interestingly, 97% of these region-specific circRNAs showed higher circularisation rate in CB, suggesting the existence of trans-factors that favour circRNA formation in CB.

**Figure 7.**
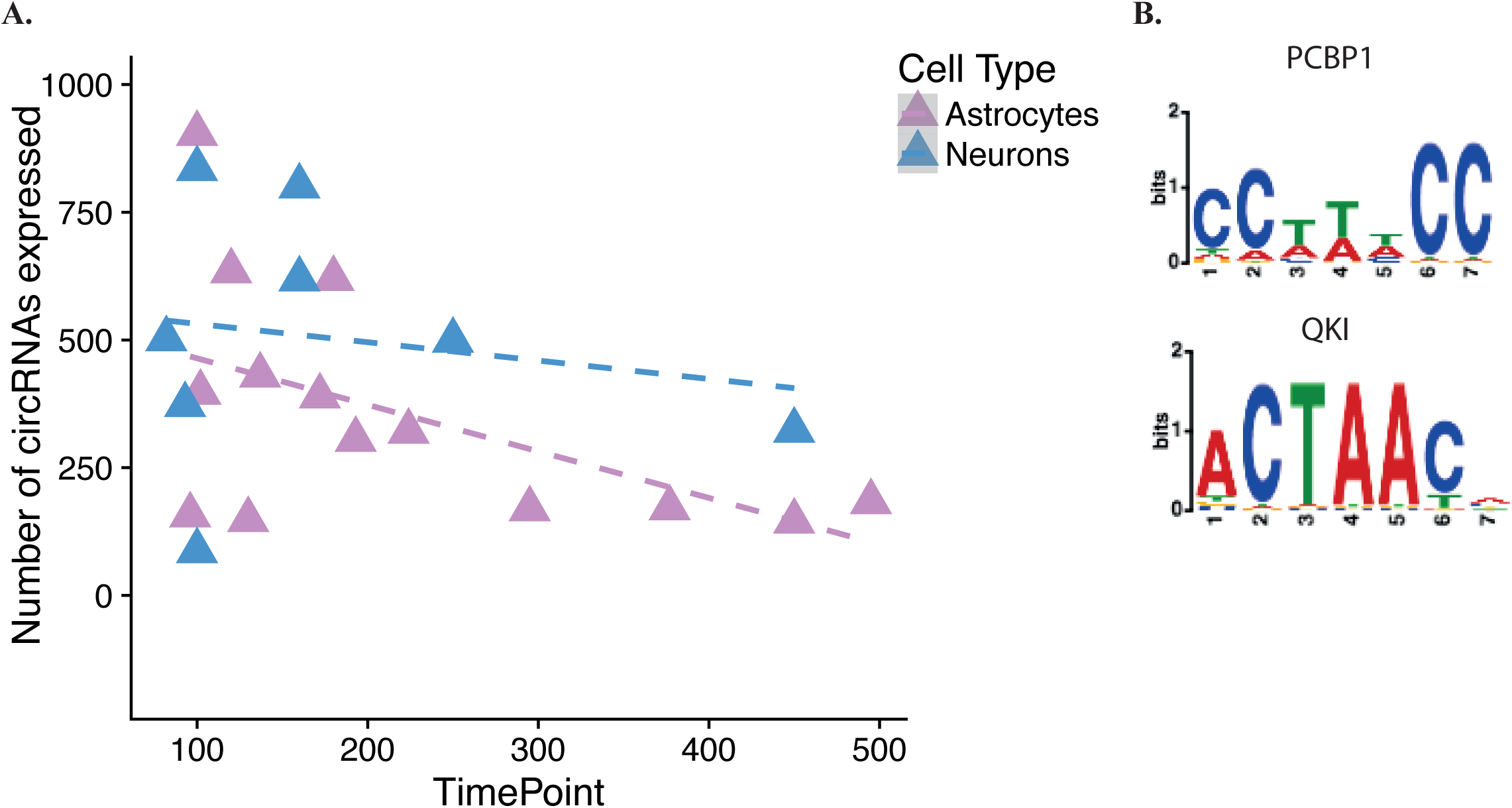
CircRNA expression changes during cellular maturation in neurons and astrocytes. **(A)** The number of circRNAs expressed in astrocytes and neurons (y-axis) vs. the maturation time point in organoid cultures (x-axis). **(B)** Top two enriched RBP motifs for neuron-specific circRNAs.

We also investigated how circRNA expression varies with age in cerebral cortex and cerebellum from control individuals. We found that the total number of circRNAs expressed showed an increasing trend with age in both brain regions, which was borderline-significant using a linear model with correction for covariates (p = 0.047; DS1 data; Figure 6c). This observation was not statistically significant in DS2, likely due to the lower sample size (p > 0.05). In addition, we assessed how individual circRNAs change in expression across the life span, using both a linear model and spline regression with inflexion points at 0, 10, 20 and 40 years. We did not identify any circRNAs that significantly changed in expression with age, after correction for co-variates and multiple testing (Methods). This is in contrast with gene expression changes with age, where as expected we identified over 500 genes significantly changing in expression across the life-span, most of which changed in the early developmental period (0–10 years), consistent with previous data [47, 48]. Our data indicate that the variation of circRNA expression with age is less pronounced than that of gene expression, with a mild increasing trend across the life span.

### Cell-type specific and developmental variation of circRNA expression

We next investigated circRNA expression in individual brain cell types by generating ribo-depleted RNA-seq data from cultured human primary astrocytes as well as *in-vitro* differentiated neurons (2-weeks differentiation; Methods). We found a much higher number of circRNAs expressed in neurons than in astrocytes: 3,601 in neurons vs. 93 in astrocytes, of which 74 were present in both cell types (Methods).

To investigate if this large difference in the number of circRNAs expressed in neurons and astrocytes is replicable, and to further investigate circRNA expression during cellular maturation, we mined RNA-seq data from human brain organoids immunopanned using either neuronal or astrocyte markers at various time points of organoid development (0–495 days) [22]. We observed variable numbers of circRNAs expressed (>=0.1 CPM) at early maturation time points (0–150 days), followed by a progressive decrease in the number of circRNAs expressed in mature neurons and astrocytes (>150 days; Figure 7a, p= 0.003, linear model t-statistic). Mature neurons indeed expressed higher number of circRNAs than mature astrocytes: 420 circRNAs were expressed in mature neurons vs. 318 in mature astrocytes (at a minimum of 0.1 CPM, in at least two mature cell samples, i.e. >150 days of maturation). However, the difference was less pronounced than what we had observed between neurons differentiated for 2 weeks and cultured astrocytes. This observation is consistent with our data showing a progressive decrease in the number of circRNAs with cellular maturation.

### Predicting trans-factors involved in circRNA formation in the human brain

Although the formation of circRNAs is strongly influenced by *cis*-factors, in particular the presence of repeats in the flanking introns [26, 49, 50], such factors cannot explain circRNA expression differences between cell types or brain regions. Among *trans-factors*, Quaking (QKI) has been identified as a regulator of circRNA formation during epithelial-to-mesenchymal transition [51], while Muscleblind (MBNL) has been shown to regulate circRNA formation from its parental gene in *Drosophila* [36]. We hypothesized that additional RNA-binding proteins may play similar roles in the human brain. We thus carried out an enrichment analysis of circRNA-flanking regions for RNA binding protein (RBP) consensus binding sites, using MEME-ChIP [52] and a 100 bp window around each circRNA end (Methods). This enrichment analysis was carried out for astrocyte-specific circRNAs and neuron-specific circRNAs, defined as circRNAs detected in one cell type by the above criteria, but not the other. QKI served as a positive control, since it was four-fold higher in expression in organoid-derived mature astrocytes vs. neurons, and it is known as a potent regulator of circRNA formation during epithelial-to-mesenchymal transition [51]. Indeed, QKI binding sites were enriched in the set of astrocyte-specific circRNAs (Supplementary Table 4), confirming the validity of our approach. In addition, astrocyte-specific circRNAs also showed enrichment for PCBP1 binding sites. Remarkably, the top enriched RBP binding sites for neuron-specific circRNAs were PCBP1 and QKI sites (Figure 7b). The enrichment of neuronal-specific circRNAs for QKI binding sites suggests that despite its lower expression levels in neurons, QKI also plays an important role in circRNA formation in these cells, acting upon neuron-specific transcripts. Our data also suggests that PCBP1, which is expressed in both cell types, may play a similarly important role in circRNA formation in both neurons and astrocytes. Three other RBPs showed binding site enrichment only for neuron-specific circRNAs: SRSF1, SRSF10 and PABPC4 (Supplementary Table 4), indicating that these RBPs may play a role in circRNA formation specifically in neurons.

### Co-expression networks identify circRNA expression differences in cerebral cortex in ASD

Previous studies [23, 43] identified gene expression differences between ASD and controls in cerebral cortex. Given the heterogeneity of ASD and the often subtle gene expression changes, advanced methods such as co-expression analyses were required to uncover gene expression changes in ASD [23, 43]. Therefore we carried out a co-expression network analysis of circRNA expression (DS1 data, CPM) using the cortex samples, after regressing out all covariates except phenotype (Methods). To address the problem of sparse data (i.e. most circRNAs being expressed in a small number of samples) we (a) used a similarity measure robust to outliers (biweight midcorrelation [53]), and (b) only included in the network analyses circRNAs expressed in at least a half of the 92 CTX samples (1,280 circRNAs). We identified 5 co-expression modules, of which one module (M4) showed significant eigengene differences between ASD and controls, with higher levels in ASD (p=0.006, Wilcoxon rank-sum test, Bonferroni corrected; Figure 8a; Methods). M4 contained 98 circRNAs (Supplementary Table 5). Interestingly, the hub of this co-expression module is a circRNA expressed from *ZKSCAN1*, which plays a role in cell proliferation and migration in hepatic cells but its role has not yet been investigated in the brain (Figure 8b). M4 also included *circHIPK3*, a circRNA implicated in cell proliferation and migration through a miRNA sponge mechanism (Figure 8b) [54, 55]. Consistent with previous data on gene expression [23], we did not identify any ASD-associated circRNA co-expression module in the cerebellum.

## DISCUSSION

The data presented here represents the first large-scale assessment of circRNA expression in the human brain, bringing further insight into an additional layer of brain transcriptome complexity.

The circRNAs included in our resource were detected in a minimum of 5 samples in DS1 and 5 samples in DS2, with >90% reproducibility of detection between the two datasets, therefore representing a set of circRNAs reliably detected in the human brain. We focussed on identifying circRNAs reproducibly expressed, rather than cataloguing very rare back-splicing events.

CircRNAs, similarly to lncRNAs, were expressed in a very specific manner, and thus observed only in a subset of samples. Despite the fact that the expression of most circRNAs was restricted to a subset of samples, we observed a remarkably high correlation between DS1 and DS2 circRNA expression levels, suggesting that the circularization rate is an intrinsic property of a given back-splice junction. To facilitate further mining of this rich resource for a wide range of biological questions, for each circRNA we provide information on the number of samples it was detected in, as well as the mean expression level in each brain region of DS1 and DS2 (Supplementary Table 2). CircRNA expression values normalized to sequencing depth for all DS1 and DS2 circRNAs are provided in Supplementary Table 2 and can be accessed as a genome browser track at: http://www.voineagulab.unsw.edu.au/circ_rna. We also provide the first resource of circRNA expression in human brain organoid cultures (Supplementary Table 6).

The analysis of this rich resource revealed several novel insights into circRNA expression in the human brain. We found that circRNA expression is characterised by major isoforms, adding a novel aspect to the notion that circRNA expression is regulated in the human brain. As more data on circRNA expression in other tissues becomes available, it will be interesting to determine whether the major isoform circRNA expression is tissue-specific, as it is often the case for mRNAs, and how this relates to the tissue specificity of alternative splicing isoforms.

We observed a significant association between AS events and circRNA expression, which is consistent with previous data from human fibroblasts [49]. The number of AS events per gene explained ~10% of the variance in the number of circRNAs produced. This indicates that AS is a relevant factor in circRNA biogenesis in the brain, but despite the complexity of AS in the brain, it only partially accounts for circRNA formation. We also found that some of the circRNA hotspot genes, expressing more than five circRNA isoforms, did not show evidence of AS. These data suggest that the interplay between circRNA formation and AS is locus-specific, in agreement with the notion that *cis*-factors, such as the presence of complementary repeat sequences, play an important role in circRNA formation [26, 49, 50]. CircRNAs can be themselves alternatively-spliced, and while we don’t address this aspect (it would require higher read length and sequencing depth [56]), an interesting future avenue is to investigate whether the complexity of circRNA AS in the human brain parallels that of linear RNAs.

The circRNA expression conservation rate between human and mouse determined in our study was consistent with that reported previously (Rybak-Wolf et al. 2015, [6]), at 8–9%. This estimate is very conservative since we (as well as Rybak-Wolf et al. 2015) required precise matching of human and mouse back-splice junction coordinates. The conservation of circRNA expression from mouse to human using the same approach is much higher, close to 30% [6], indicative of higher complexity of circRNA expression in the human brain.

The analysis of circRNA expression changes with age showed a mild increase in the total number of circRNAs expressed across the life-span (0 to 60 years; DS1). Unlike gene expression, which shows major changes in the early postnatal period, followed by a plateau after 20 years of age [47], circRNAs showed a mild increasing trend throughout the life span. Previous studies have reported that circRNAs accumulate with age in mouse and *drosophila* brain [8, 57]. The comparison of 1 month and 22 month-old mice identified more up-regulated than down-regulated circRNAs in CTX and hippocampus, at p<0.05 using a *t*-test without correction for multiple testing [57]. Given the weak statistical significance and the fact that down-regulation might be more difficult to detect for these already lowly expressed molecules, further data would be required to determine whether circRNA expression changes with age in a similar manner in mouse and human brain. In *Drosophila* CNS, the total number of circRNAs detected increases with age, while individual circRNAs only passed a permissive statistical threshold (p < 0.05, without multiple testing correction) [8]. Taken together, the, available data across species support the notion of a progressive increase in circRNA levels with age in the brain, but this effect appears to be of a low magnitude, and therefore challenging to detect at genome-wide significance.

In contrast to age-dependent changes, circRNA expression differences between CTX and CB were pronounced, overriding inter-individual variability. We identified over two hundred circRNAs differentially expressed between CTX and CB, in both DS1 and DS2 and after correction for cellular composition. We also found that more than 90% of differentially expressed circRNAs showed higher expression in CB. This is in agreement with a previous observation based on two frontal cortex and two cerebellum replicates [6], showing overall higher circRNA expression in the cerebellum. The initial observation was interpreted as a reflection of higher proportion of neurons in cerebellum than in cerebral cortex, a known property of this brain region. Here we demonstrate that the enrichment of circRNAs in the cerebellum is independent of cell-type composition.

In human brain organoids, the number of circRNAs expressed showed a progressive decrease with cellular maturation in both neurons and astrocytes. Despite the documented increase in circRNA expression during early neuronal differentiation [6, 7, 9, 10], we found that neuronal maturation beyond 150 days in organoid culture leads to an overall decrease in circRNA expression. Using data from neuronal and glial cells at the same maturation stage we found an enrichment of astrocyte-specific circRNAs for QKI binding sites, consistent with its known role in circRNA formation. We also uncovered PCBP1 as a novel candidate for circRNA expression regulation in astrocytes and neurons, and SRSF1, SRSF10 and PABPC4 as potential regulators of circRNAs in neurons, thereby highlighting these proteins as valuable candidates for further functional studies.

Finally, using a co-expression network approach we identified a circRNA co-expression module showing increased expression in ASD, the first identification of circRNA expression changes in this disorder.

Taken together our data is the first to provide information on circRNAs detected in human brain across multiple individuals. It represents a rich resource on circRNA expression in the human brain, available in a browsable format at http://www.voineagulab.unsw.edu.au/circ_rna

## METHODS

### RNA samples and RNA-seq data

The RNA-seq data published by Parikshak et al. [23] (ribo-depleted, unstranded, 50 bp paired-end), was kindly provided by Prof. Daniel Geschwind before publication. Frontal cortex samples had been obtained from Brodmann area (BA) 9, temporal cortex samples from BA 41/42 and 22, and cerebellar samples had been obtained from cerebellar vermis. We only included samples with RIN> 5 (Supplementary Table 1).

The distribution of samples between the two datasets (DS1 and DS2) is listed in Supplementary Table 1, and was based on the order in which we obtained the data. This in turn was based on the order in which RNA-seq data was generated, which was randomised across groups (Figure 1a).

RNA-seq data for benchmarking circRNA detection was generated for 5 DS1 tissue samples: 5115_ba9, 5278_ba9, 5297_ba41–42, 5308_ba41–42, 5309_ba41–42. Brain tissue from the same individuals and brain region had been obtained by our lab from the NICHD brain and tissue bank. Total RNA was extracted from ~ 100 mg of brain tissue using a Qiagen miRNeasy kit according to the manufacturer’s protocol, and treated with 1 μl DNase I (Thermo Fisher Scientific, #AM2238) per 10 μg of RNA. Each RNA sample was divided in two, for library preparation with either the TruSeq Stranded Total RNA Ribozero kit or the TruSeq Stranded mRNA kit (which includes polyA selection). Library preparation was carried out at the UNSW Ramaciotti Centre for Genomics, followed by sequencing on an Illumina NextSeq 500 sequencer to obtain 100 bp paired-end reads (Supplementary Table 1).

For generating RNA-seq data from astrocytes and neurons, total RNA was extracted from human primary astrocytes and from neurons derived from human fetal neural progenitors.

Human primary astrocytes (Lonza, #CC-2565) stably expressing GFP from pCMV6-AC-GFP had been generated by selection with G418 (Thermo Fisher Scientific, #10231027) at 800μg/ml. Cells were cultured in RPMI GlutaMAX™ (Thermo Fisher Scientific, #35050061) supplemented with 10% foetal bovine serum, 1% streptomycin (10,000 μg/ml), 1% penicillin (10,000 units/ml) and 1% Fungizone (2.5 μg/ml) and seeded into 6-well tissue culture plates at a density of 0.5 × 10^6^ cells 24 hours prior to RNA extraction. Total RNA was extracted using TRIzol^®^ reagent and a Qiagen miRNeasy kit and treated with 1 μl DNase I (Thermo Fisher Scientific, #AM2238) per 10 μg of RNA.

Neuronal differentiation of human neural progenitors stably transfected with pLRC-GFP was carried out for 2 weeks as previously described [58]. RNA extraction was carried out using a Qiagen miRNeasy kit, with on-column DNase digestion [58].

RNA samples from astrocytes (n=1), neurons (n=1) were depleted of ribosomal RNA using the Epicentre Ribo-zero kit, according to the manufacturer’s protocol. Library preparation using the Illumina TruSeq Stranded kit (http://www.illumina.com/products/truseq_stranded_total_rna_library_prep_kit.html) and sequencing on a NextSeq 500 Illumina sequencer were carried out at the UNSW Ramaciotti Centre for Genomics, generating 75 bp paired-end reads (Supplementary Table 1).

The RNA-seq data from human brain organoids was downloaded from SRA, accession number GSE99951.

### CircRNA benchmarking analysis

For five brain tissue samples (5115_ba9, 5278_ba9, 5297_ba41–42, 5308_ba41–42, 5309_ba41–42), the polyA+ and ribo-depleted stranded data generated in the present study, as well as the ribo-depleted unstranded data from Parikshak et al. were mapped to the human genome (hg19) using STAR [59], and the chimeric read alignments were used as input for either DCC [27] or CIRCexplorer [26], run with default parameters. CircRNAs were filtered to include those detected by at least 2 back-spliced junction reads in a minimum of 2 distinct samples. CircRNA counts were normalised to library size to obtain counts-per-million (CPM).

### Human brain circRNA dataset

RNA sequencing reads from DS1 and DS2 brain samples were aligned to the human genome (hg19) using STAR [59] as described above. Gene-level expression was assessed with *featureCounts*, as implemented in STAR. Samples with more than 3 standard deviations away from the mean of mean inter-sample correlations, as well as outliers on PCA analysis were eliminated from further analyses. For each dataset (DS1 and DS2), genes were filtered for expression at a minimum of 1 RPKM in at least 30% of the smallest sample group within each dataset (i.e. 6 samples in DS1 and 4 samples in DS2). CircRNAs were identified using STAR’s chimeric read alignments as input for DCC [27], with default parameters for paired-end un-stranded data. Counts for circRNAs with identical genomic coordinates on opposite strands were summed, given the unstranded nature of the data. CircRNA counts were normalised to the total number of uniquely aligned reads, to obtain counts per million (CPM). For each circRNA, its corresponding linear junction counts were quantified using splice junction counts generated by STAR. The downstream and upstream linear junction counts were calculated as the sum of all linear junction reads spanning the start- and the end-circRNA coordinate respectively (schematic representation below). Linear junction reads were also normalised to library size (i.e. the total number of uniquely aligned reads).

**Figure 8.**
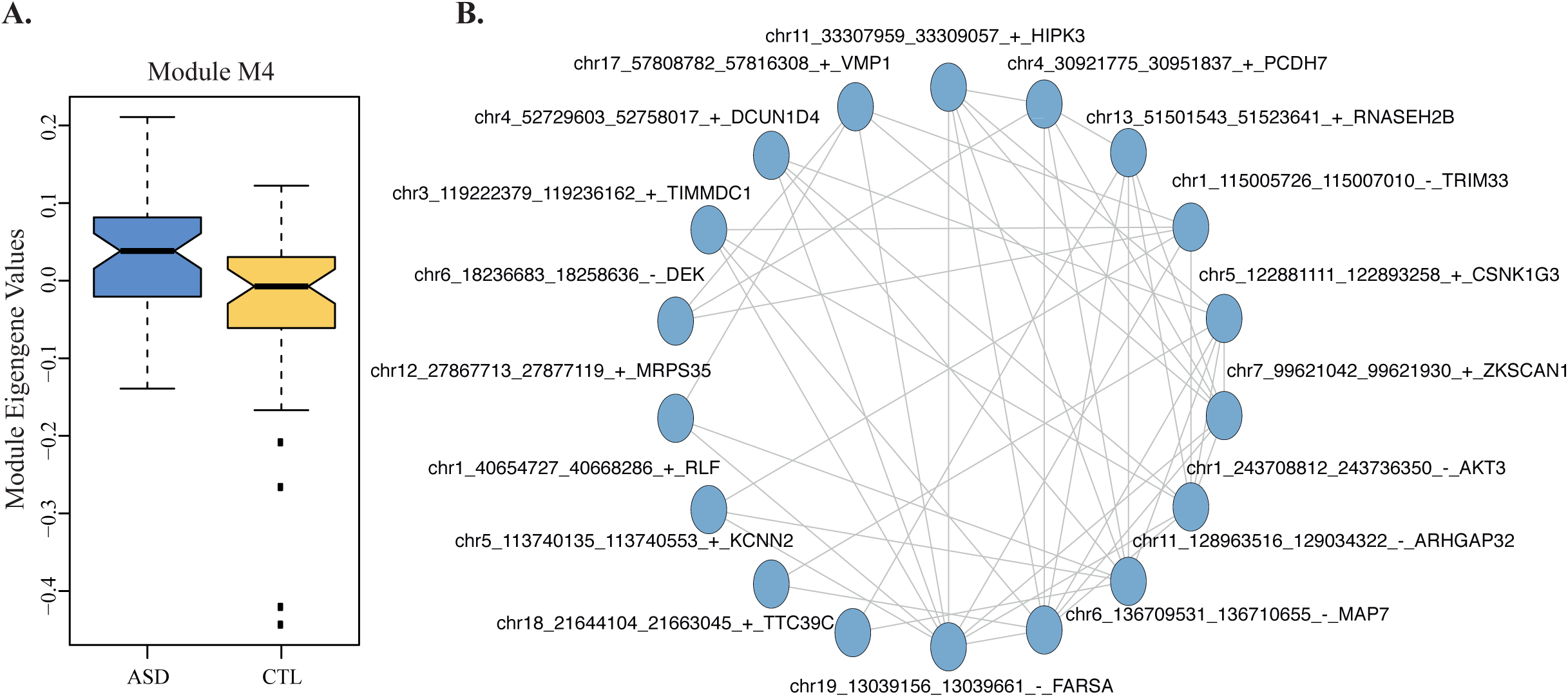
Co-expression networks identify circRNA expression changes in ASD. **(A)** Module eigengene values for M4 show significant differences between ASD and control CTX samples. p=0.006, Wilcoxon rank-sum test, Bonferroni corrected. Boxplots were generated using the boxplot function in R; the horizontal line represents the median, boxes extend between the first and third quartiles, and whiskers extend to 1.5 IQR (inter-quartile range) from the box. Notches mark +/-1.58 IQR/sqrt(n), where n represents the number of data points. **(B)** Network plot of M4, showing the top 20 circRNAs by kME (blue circles), and the top 50 connections between them as edges.

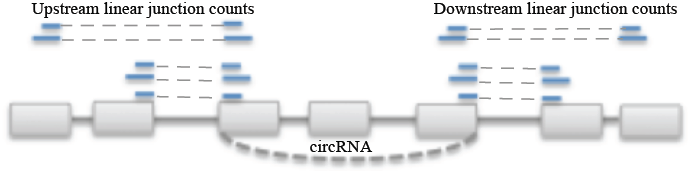

#### CircRNA annotation relative to genomic features

Since the RNA-seq data was un-stranded, circRNAs were assigned a strand based on their overlap with exon junctions as follows: if either the start or the end overlapped an annotated exon junction, the circRNA was assigned the strand of the corresponding transcript. If neither start nor end overlapped an exon junction, or if they overlapped exon junctions on both strands, the strand was set as ambiguous. CircRNAs left with ambiguous strand annotation were next overlapped with gene intervals and assigned a strand in the same manner as above. CircRNA genomic annotation was done separately for the start and end coordinates with the following hierarchy: exon junction>exonic>intronic>genic (Supplementary Figure 3). CircRNAs were annotated relative to circBase downloaded 09.2017. All genomic annotations were performed relative to Ensembl GRCh37 transcript annotations downloaded 10.2014.

#### CircRNA filtering

To select robustly expressed circRNAs we used the overlap between DS1 and DS2 as a guide for filtering criteria. We assessed the overlap between circRNAs detected in DS1 and DS2 when requiring circRNAs to be expressed at a minimum of either 2 read counts (a permissive criterion) or 0.1 CPM (stringent criterion) in a range of numbers of samples from 1 to 10 (Supplementary Figure 2e). We found that the overlap between DS1 and DS2 increased with the number of samples we required expression in, as expected, and plateaued at 5 samples. Notably, when requiring expression in a minimum of 5 samples, the overlap between DS1 and DS2 was above 90% using either the permissive or the stringent criteria, and thus we used this filtering parameter for inclusion of circRNAs in further analyses. However, we also include in the supplementary information (Supplementary Table 2) the number of samples in which each circRNAs is expressed at a higher expression threshold (>= 0.1 CPM), to allow this resource to be easily used to identify highly expressed circRNAs.

##### Gene ontology enrichment analyses of circRNA-producing genes

GO enrichment analysis was carried out using the intersection of circRNA producing genes from DS1 and DS2, and the intersection of all genes expressed in DS1 and DS2 as background. The enrichment analysis was done using GOseq [60], with correction for gene length and Benjamini and Hochberg (BH) correction for multiple testing.

##### Conservation of circRNA expression

To assess circRNA expression conservation between human and mouse brain, we first converted hg19 human genomic coordinates to mm9 mouse coordinates using the liftOver tool from the UCSC genome browser. The converted coordinates were then interrogated using the mouse circRNA expression data from Rybak Wolf et al. [6] (Supplementary Table S1 of that study). The % conservation was calculated as the percentage of human circRNAs in DS1 and DS2 respectively, for which their converted genomic coordinates were found as circRNAs expressed in the mouse brain.

##### CircRNA flanking-intron sequence complementarity analysis

Introns flanking circRNA back-splice junctions were used as input for autoBLAST [39], which employs BLAST with the following settings: parameters: blastn, word size 7, output format 5, to determine sequence complementarity in circRNA flanking introns. Intron-pairing score for a given circRNA was defined as the number of reverse-complementary matches with a minimum bit score of 20.

##### Alternative splicing analysis

rMATS [40] was run with default parameters for 50 bp paired-end reads on DS1 samples contrasting CTX (Sample 1) and CB (Sample 2), and identified 145,995 cassette exons. Alternatively spliced exons were then filtered within each brain region, to include those supported by at least 2 inclusion junction reads and 2 skipping junction reads in two independent samples, leading to 33,207 AS exons in CTX and 24,380 AS exons in CB. These data are consistent with recent alternative splicing analyses in the human brain: Takata et al. 2017 identified 29,271 alternatively spliced exons using RNA-seq data from the Common Mind consortium (206 human dorsolateral-prefrontal cortex samples) [61].

All comparisons between circRNA expression and AS were carried out in control samples.

##### *In-silico* deconvolution

To estimate *in-silico* the proportion of individual cell types in brain tissue samples, we used DeconRNAseq, which is specifically designed for RNA-seq data [45], and two distinct reference transcriptome datasets: (a) CAGE data from cultured neurons and astrocytes from the FANTOM5 consortium [62], and (b) RNA-seq data from immunopanned human astrocytes and neurons from the Barres lab (Zhang et al., 2016 [46]). We found a very high correlation between the neuronal proportion estimates based on the two reference transcriptomes (Spearman *rho* > 0.9, Supplementary Figure 8a). We also found high correlation between neuronal proportion estimates obtained by FANTOM5 or Zhang et al., 2016 reference transcriptomes and the expression level of neuronal-specific genes *(MAP2, RBFOX1*, Supplementary Figure 8b). Given the highly similar results obtained with neuronal proportion estimates based on the two reference transcriptome datasets, we used the cell-type reference transcriptome data from Zhang et al., 2016 in all downstream analyses. However, to show that the significant changes in circRNA expression between brain regions is robust to the neuronal proportion estimates, we also report the differential expression results obtained with FANTOM5 reference-based estimates in Supplementary Table 7.

##### Differential expression with brain region and age

To assess differential circRNA expression with brain region and age, we applied a linear model to circRNA expression levels normalised to their linear transcript (CI), using the control samples (DS1). Only circRNAs expressed in at least half of the samples were included in the analysis. Data was log2-transformed with an offset of 0.5. The following variables were included in the model: proportion of neurons, sex, RNA integrity number, sequencing batch, brain bank, age and brain region (CTX and CB). The same linear model was applied to gene-level expression data (RPKM). P-values for the linear model, obtained using the *summary* function in R, were corrected for multiple testing using a BH correction as implemented in the *multtest* Bioconductor package (https://www.bioconductor.org/packages/release/bioc/html/multtest.html). We did not identify any circRNAs differentially expressed with age (adjusted p < 0.05). We also assessed circRNA expression changes with age using spline regression, with knots at 0, 10, 20, and 40 years old, and degree=1, using the basic spline *(bs)* function in the *splines* R package (https://cran.r-project.org/web/packages/splines/index.html), with the same result. The result remained the same, whether or not we included the proportion of neurons in the model.

501 circRNAs were differentially expressed between brain regions in DS1. To assess whether this result is replicable in the smaller dataset (DS2), we applied the same data analysis approach as above in DS2. We included both ASD and control DS2 samples, in order to increase statistical power, and added phenotype to the list of co-variates. 266 circRNAs were replicated in DS2 and are listed in Supplementary Table 7.

##### Co-expression network analyses

Network analysis was carried out in the larger dataset using circRNAs expressed in at least half of the CTX samples (DS1) using the blockwiseModules function in the WGCNA R package [63] with the following parameters: power =10, networkType = "signed", corFnc="bicor", minModuleSize=10, mergeCutHeight=0.15. The beta power was chosen so that the network fulfilled scale-free topology (r^2^ > 0.8). CircRNAs were assigned to a module based on their correlation to the module eigengene value (kME > 0.1) and a significant BH-corrected p-value for this correlation (adjusted p < 0.05). kME values are listed in Supplementary Table 5.

##### RT-PCR validation of circRNAs

22 circRNAs present in more than a third of the brain tissue samples were chosen for validation. RT-PCR was performed with divergent primers, on three independent RNA samples. RNA was extracted using a Qiagen miRNeasy kit, with on-column DNase digest. PCR amplification was carried out using BioRad iTaq Polymerase for 35 cycles, on an ABI ViiA7 cycler. Primer sequences for RT-PCR experiments are listed in Supplementary Table 8.

### Astrocyte and neuron circRNA dataset

Mapping and circRNA quantification for RNA-seq data generated in the present study (cultured neurons and astrocytes), and data from brain organoids were carried out as described above for brain tissue data, with the difference that the RNA-seq data generated in this study is strand-specific, which was specified as a parameter in DCC. Gene-level expression data was filtered to include genes expressed at a minimum of 1 RPKM in at least 1 sample in each dataset. CircRNAs were included in the dataset if they were expressed at a minimum of 0.1 CPM in at least 1 sample.

CircRNAs expressed in neurons and astrocytes in the brain organoid data were defined as circRNAs expressed at >=0.1 CPM in at least two samples of the fully matured neuronal and astrocyte samples (> 150 days) respectively. Neuron-specific and astrocyte-specific circRNAs were defined as circRNAs fulfilling the above criteria in one cell type but not the other.

#### RBP binding site enrichment analysis

DNA sequence corresponding to a window of 100 bp upstream and 100 bp downstream of the start and end coordinates of each circRNA was retrieved using the *fastaFromBed* function in BedTools [64]. For each set of circRNAs (i.e. astrocyte-specific and neuron-specific) both the start and end sequences were included in the RBP enrichment analysis. Local enrichment analysis was carried out using CENTRIMO, implemented in MEME-ChIP. MEME-ChIP was run on the online server (http://meme-suite.org/tools/meme-chip) with default parameters, using the “RNA, DNA encoded option”, and the RBP binding site data from Ray et al. 2013 [65], *Homo sapiens*. The analysis was limited to the strand corresponding to the provided sequence, as recommended for RNA data. The background set of sequences consisted of 100 bp windows around the start and end coordinates of 100,000 exons randomly selected from genes not forming circRNAs in the brain or in the organoid culture data.

## ACKNOWLEDGEMENTS

The authors would like to thank Dr. Cristopher Pardy and Dr. Jim Fang for technical support in the initial stages of the project. This work was supported by an NHMRC project grant and an ARC Future fellowship to IV.

## DATA AVAILABILITY

Sequencing data from this study is available in SRA [ID:TBD] before publication.

## AUTHOR CONTRIBUTIONS

IV conceived the study and supervised all aspects of the project. AG, FA, and IV analysed data, AG carried out experimental validations, AG and IV wrote the manuscript.

## COMPETING INTERESTS

The authors declare no competing financial interests.

## SUPPLEMENTARY FIGURES

**Supplementary Figure 1. Characterisation of DS1 and DS2 sample composition.** **(A)** DS1. **(B)** DS2. p: Wilcoxon rank-sum test p-values for age, and Fisher test p-values for gender ratios. Boxplots were generated using the boxplot function in R; the horizontal line represents the median, boxes extend between the first and third quartiles, and whiskers extend to 1.5 IQR (inter-quartile range) from the box. Notches mark +/-1.58 IQR/sqrt(n), where n represents the number of data points.

**Supplementary Figure 2. Benchmarking of circRNA expression data**. **(A)** Barplots of false positive rates defined as the % of circRNAs detected in each DS1 sample that were also detected in polyA+ data from the same sample, normalized for the sequencing depth. **(B)** Barplots of sequencing depth (millions of uniquely mapped reads) for DS1 libraries, as well as RNA-seq data generated in the present study (polyA+ and RD_stranded). RD: ribo-depleted. **(C)** Barplots of the number of circRNAs detected in polyA+ libraries (i.e. false-positives) for DS1 data, ribo-depleted stranded data, as well as those common between both types of data. RD: ribo-depleted. Left: DCC, Right: CIRCexplorer. **(D)** Scatterplots of normalized circRNA expression levels (CPM: counts per million) quantified by DCC and CIRCexplorer. *r*: Pearson correlation coefficient. **(E)** CircRNA detection using DCC across various expression thresholds. Number of circRNAs detected by using a threshold of either 2 back-splice junction reads (left), or 0.1 CPM (right), in a minimum of 1 to 10 independent samples.

**Supplementary Figure 3. CircRNA annotation relative to genomic features**. exonJ-exonJ: circRNAs for which both ends correspond to annotated exon junctions. The rest of circRNAs were annotated based on the overlap of at least one end with genomic features, using the following hierarchy: exonJ > exonic > intronic > intergenic.

**Supplementary Figure 4. Comparison of circRNA expression and linear gene expression in CTL samples**. **(A)** and **(B)**. Scatterplots for DS1 **(A)** and DS2 **(B)** displaying circular junction expression vs. the expression of the corresponding linear junctions, quantified as the maximum of the upstream and downstream junctions (left panel), circular junction expression vs. parental gene expression (centre), and linear junction expression for circRNA-forming junctions vs. parental gene expression (right panel) **(C)**. Boxplots displaying the mean expression of circRNA-genes and non-circRNA forming genes in DS1 (left) and DS2 (right). Boxplots were generated using the boxplot function in R; the horizontal line represents the median, boxes extend between the first and third quartiles, and whiskers extend to 1.5 IQR (inter-quartile range) from the box. Notches mark +/-1.58 IQR/sqrt(n), where n represents the number of data points.

**Supplementary Figure 5. Intron-pairing rank of circRNA major isoforms.** Genes are classified based on how many circRNAs they express, and for each class (X-axis), the intron pairing rank of the major isoform is plotted in % (Y-axis). Rank=1: highest intron pairing score; Rank=10: lowest intron pairing score. Only genes expressing up to 10 circRNAs are plotted.

**Supplementary Figure 6. CircRNA expression variability**. **(A)** Scatterplots of mean expression (x-axis) and standard deviation (y-axis). Left: genes, centre: splice junctions, right: circRNAs. CV: mean coefficient of variation. **(B)** Histograms displaying the number of samples in which genes (left), splice junctions (centre), and circRNAs (right) were expressed.

**Supplementary Figure 7. Estimated proportion of neurons in DS1 and DS2**. (**A)** Boxplots displaying the estimated proportion of neurons across brain regions and phenotypes, in DS1 (left) and DS2 (right). Boxplots were generated using the geom_boxplot and geom_violin functions in R; the horizontal line represents the median, boxes extend between the first and third quartiles, and whiskers extend to 1.5 IQR (inter-quartile range) from the box. Notches mark +/-1.58 IQR/sqrt(n), where n represents the number of data points. (**B)** Scatterplots of first principal component values of gene expression data (PC1, y-axis) vs. estimated proportion of neurons. (**C)** Scatterplots of first principal component values of circRNA expression data (PC1, y-axis) vs. estimated proportion of neurons. All neuronal proportion estimates are based on reference transcriptome data from Zhang et al. 2016.

**Supplementary Figure 8. Assessment of cellular composition estimates**. **(A)** Scatterplot of estimated neuronal proportions based on the FANTOM5 and Zhang et al. reference transcriptome data in DS1 and DS2; *rho:* Spearman correlation coefficient. **(B)** Scatterplot of gene expression levels of two neuronal-specific genes *(RBFOX1-top* row, and MAP2-bottom row; y-axis) vs. estimated neuronal proportions (x-axis). *rho:* Spearman correlation coefficient.

## SUPPLEMENTARY TABLES

**Supplementary Table 1.** Sample description (DS1 and DS2) and summary of mapping results for all RNA-seq data included in this study.

**Supplementary Table 2.** CircRNA annotation and expression data. A. DS1 circRNA annotation. B. DS2 circRNA annotation. C. DS1 circRNA expression (CPM). D. DS2 circRNA expression (CPM)

**Supplementary Table 3.** Gene Ontology enrichment of circRNA-producing genes.

**Supplementary Table 4.** MEME-ChIP results

**Supplementary Table 5.** WGCNA circRNA results. kME: correlation between individual circRNA expression and module eigengene values. pvalBH: Student p-values for the kME correlation values, corrected for multiple testing using a Benjamini and Hochberg correction.

**Supplementary Table 6.** CircRNA expression during astrocyte and neuronal maturation. Expression values are in CPM. Sample names include SRA id, day of the maturation, time point and either “Hepa” for the astrocyte immunopanned samples of “Thy-1” for neuronal immunopanned samples.

**Supplementary Table 7.** Brain region-specific circRNAs

**Supplementary Table 8.** Primer sequences for RT-PCR validation of circRNAs.

