## Supplementary_Figure_1 for "The Landscape Of Circular RNA Expression In The Human Brain"

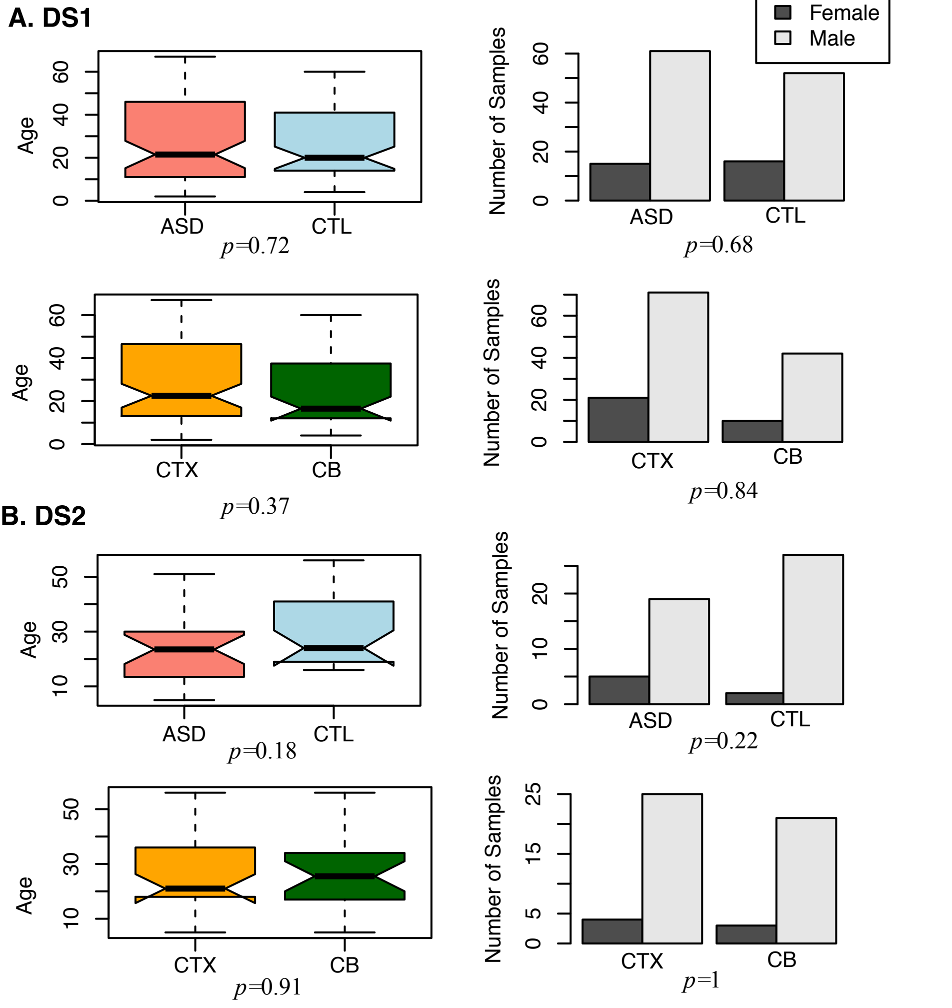


**Supplementary Figure 1. Characterisation of DS1 and DS2 sample composition. (A)** DS1. **(B)** DS2. *p:* Wilcoxon rank-sum test p-values for age, and Fisher test p-values for gender ratios. Boxplots were generated using the boxplot function in R; the horizontal line represents the median, boxes extend between the first and third quartiles, and whiskers extend to 1.5 IQR (inter-quartile range) from the box. Notches mark +/-1.58 IQR/sqrt(n), where n represents the number of data points.
