## Supplementary_Figure_2 for "The Landscape Of Circular RNA Expression In The Human Brain"

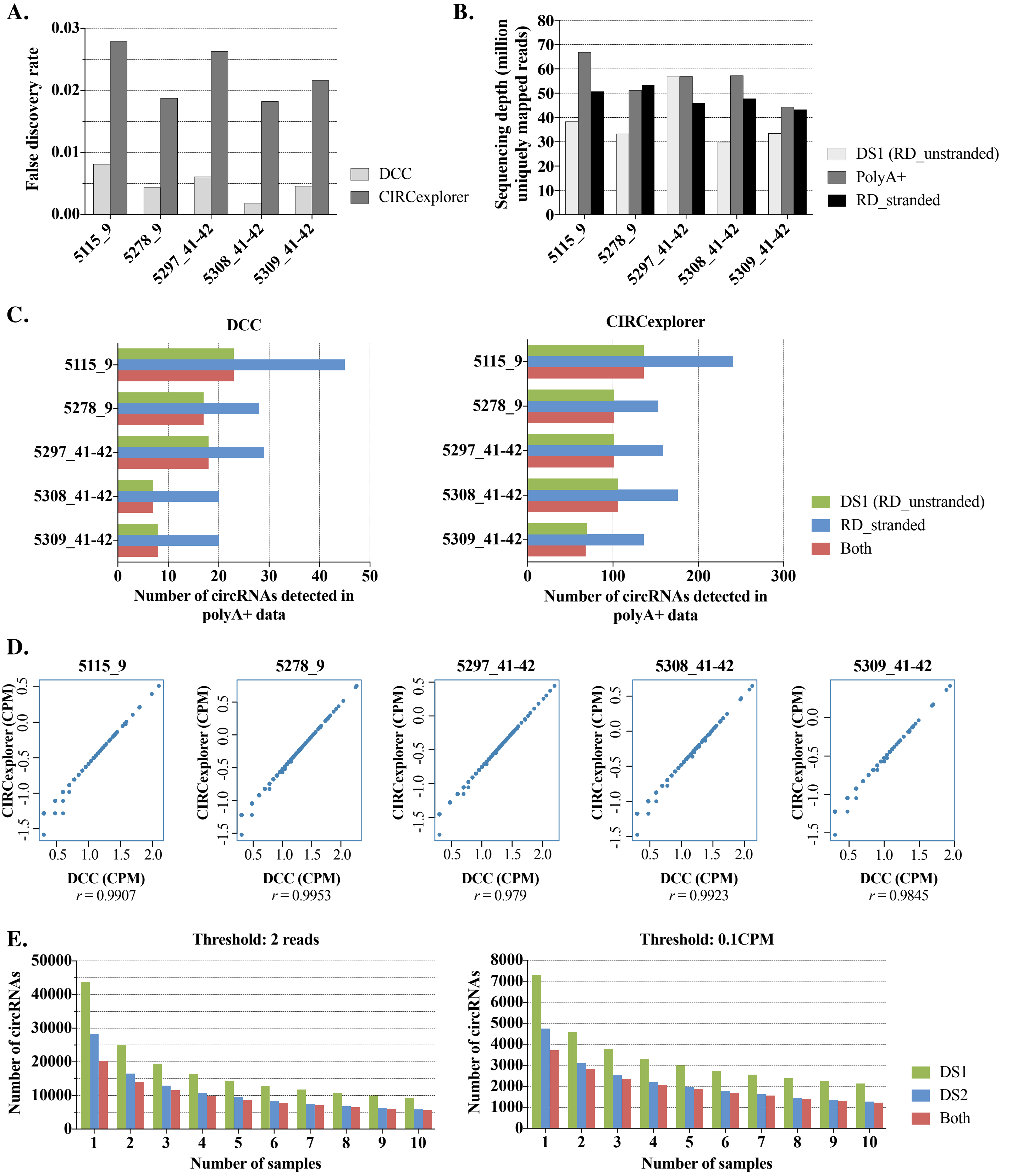


**Supplementary Figure 2. Benchmarking of circRNA expression data. (A)** Barplots of false positive rates defined as the % of circRNAs detected in each DS1 sample that were also detected in polyA+ data from the same sample, normalized for the sequencing depth. **(B)** Barplots of sequencing depth (millions of uniquely mapped reads) for DS1 libraries, as well as RNA-seq data generated in the present study (polyA+ and RD_stranded). RD: ribo-depleted. **(C)** Barplots of the number of circRNAs detected in polyA+ libraries (i.e. false-positives) for DS1 data, ribo-depleted stranded data, as well as those common between both types of data. RD: ribo-depleted. *Left:* DCC, *Right:* CIRCexplorer. **(D)** Scatterplots of normalized circRNA expression levels (CPM: counts per million) quantified by DCC and CIRCexplorer. *r* : Pearson correlation coefficient. **(E)** circRNA detection using DCC across various expression thresholds**.** Number of circRNAs detected by using a threshold of either 2 back-splice junction reads (left), or 0.1 CPM (right), in a minimum of 1 to 10 independent samples
