## Supplementary_Figure_3 for "The Landscape Of Circular RNA Expression In The Human Brain"

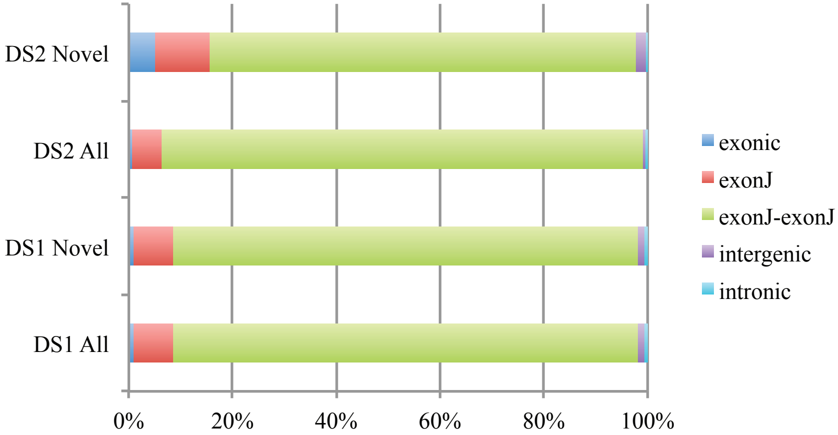


**Supplementary Figure 3. CircRNA annotation relative to genomic features.** exonJ-exonJ: circRNAs for which both ends correspond to annotated exon junctions. The rest of circRNAs were annotated based on the overlap of at least one end with genomic features, using the following hierarchy: exonJ > exonic > intronic > intergenic.
