## Supplementary_Figure_4 for "The Landscape Of Circular RNA Expression In The Human Brain"

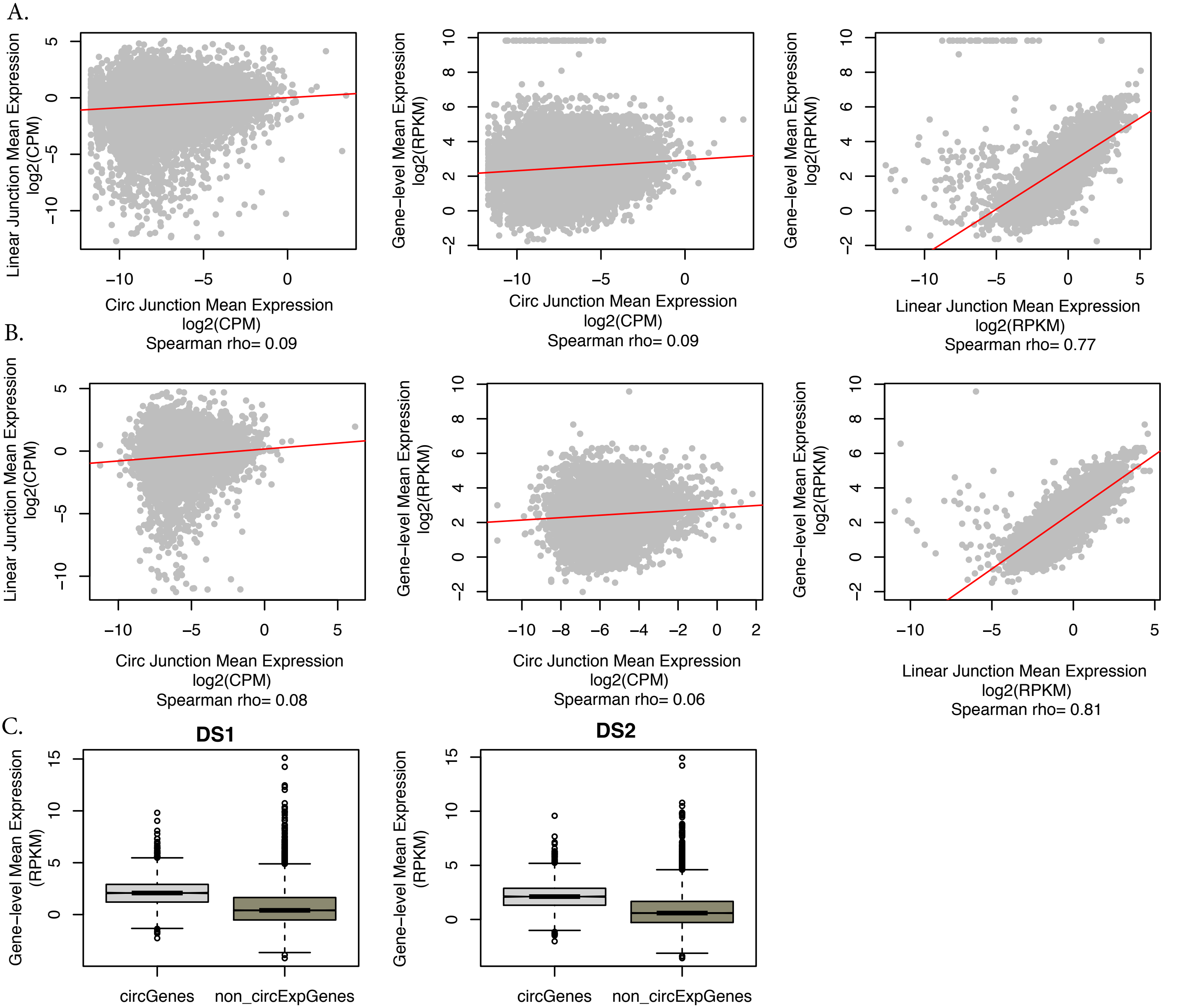


**Supplementary Figure 4. Comparison of circRNA expression and linear gene expression in CTL samples. (A)** and **(B)**. Scatterplots for DS1 (A) and DS2 (B) displaying circular junction expression vs. the expression of the corresponding linear junctions, quantified as the maximum of the upstream and downstream junctions (left panel), circular junction expression vs. parental gene expression (centre), and linear junction expression for circRNA-forming junctions vs. parental gene expression (right panel). **(C)**. Boxplots displaying the mean expression of circRNA-genes and non-circRNA forming genes in DS1 (left) and DS2 (right). Boxplots were generated using the boxplot function in R; the horizontal line represents the median, boxes extend between the first and third quartiles, and whiskers extend to 1.5 IQR (inter-quartile range) from the box. Notches mark +/-1.58 IQR/sqrt(n), where n represents the number of data points.
