## Supplementary_Figure_5 for "The Landscape Of Circular RNA Expression In The Human Brain"

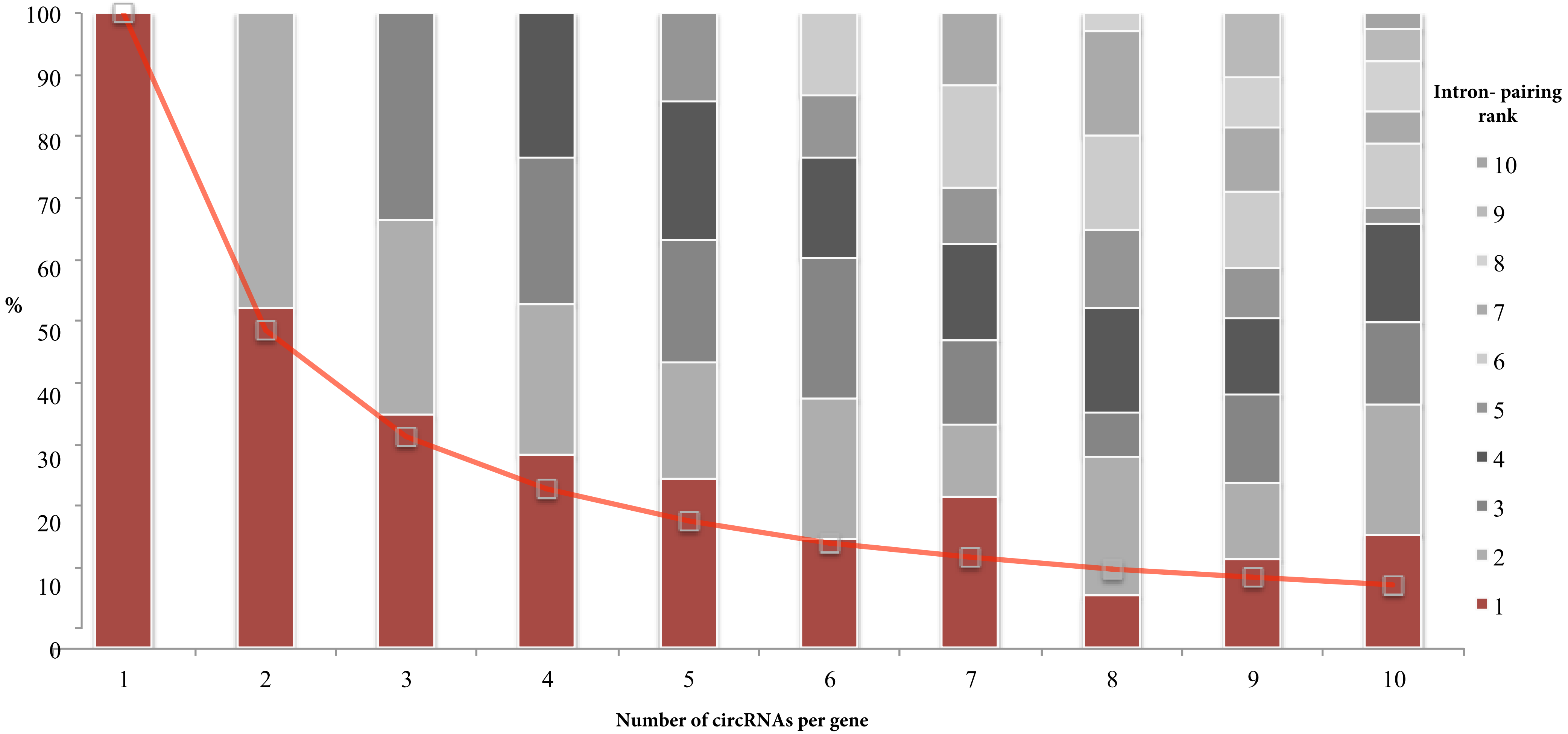


**Supplementary Figure 5. Intron-pairing rank of circRNA major isoforms.** Genes are classified based on how many circRNAs they express, and for each class (X-axis) the intron-pairing rank of the major isoform is plotted in % (Y-axis). Rank=1: highest intron pairing score; Rank=10: lowest intron pairing score. Only genes expressing up to 10 circRNAs are plotted.
