## Supplementary_Figure_6 for "The Landscape Of Circular RNA Expression In The Human Brain"

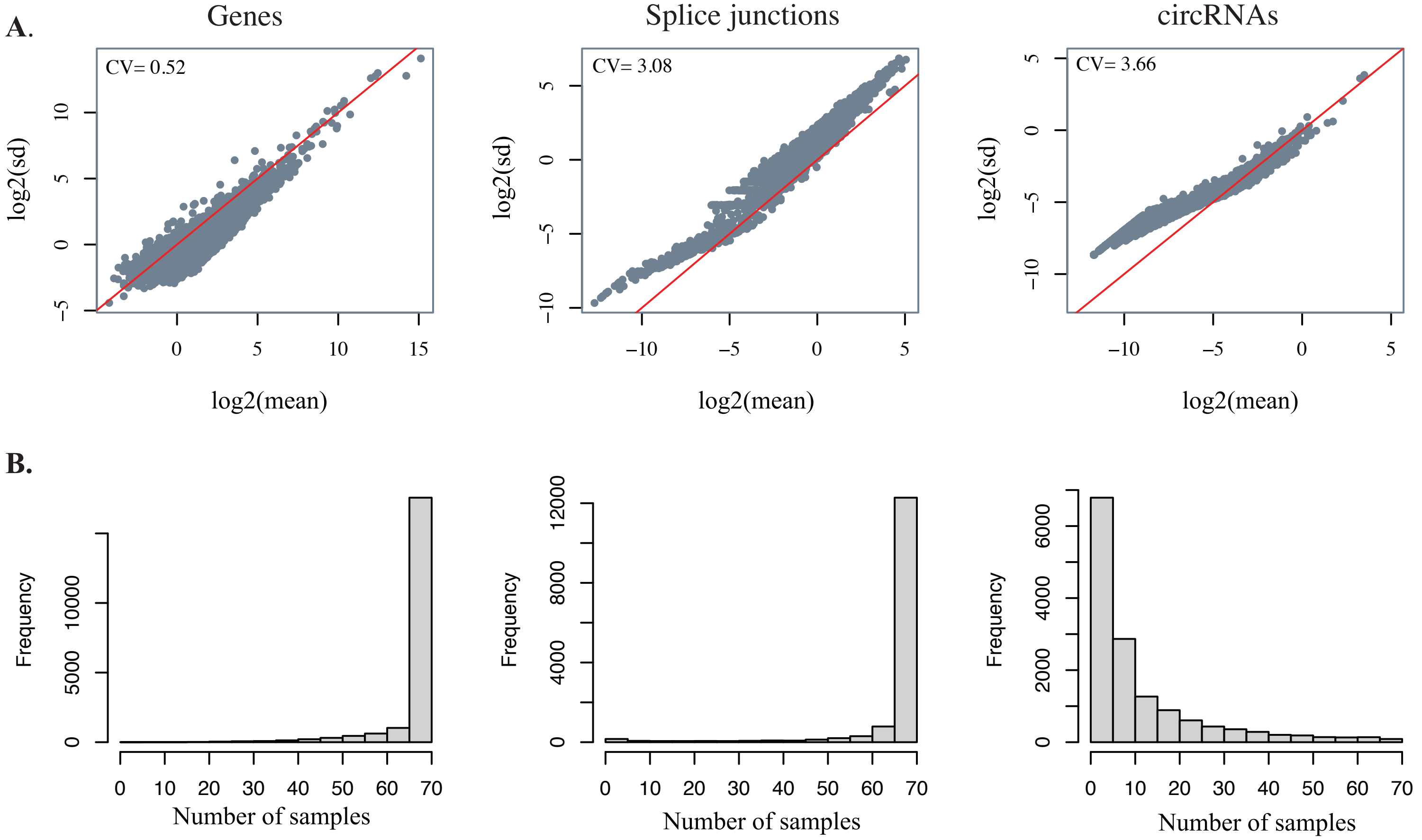


**Supplementary Figure 6. CircRNA expression variability. (A)** Scatterplots of mean expression (x-axis) and standard deviation (y-axis). *Left*: genes, *centre*: splice junctions, *right*: circRNAs. CV: mean coefficient of variation. **(B)** Histograms displaying the number of samples in which genes (*left*), splice junctions (*centre*), and circRNAs (*right*) were expressed.
