## Supplementary_Figure_7 for "The Landscape Of Circular RNA Expression In The Human Brain"

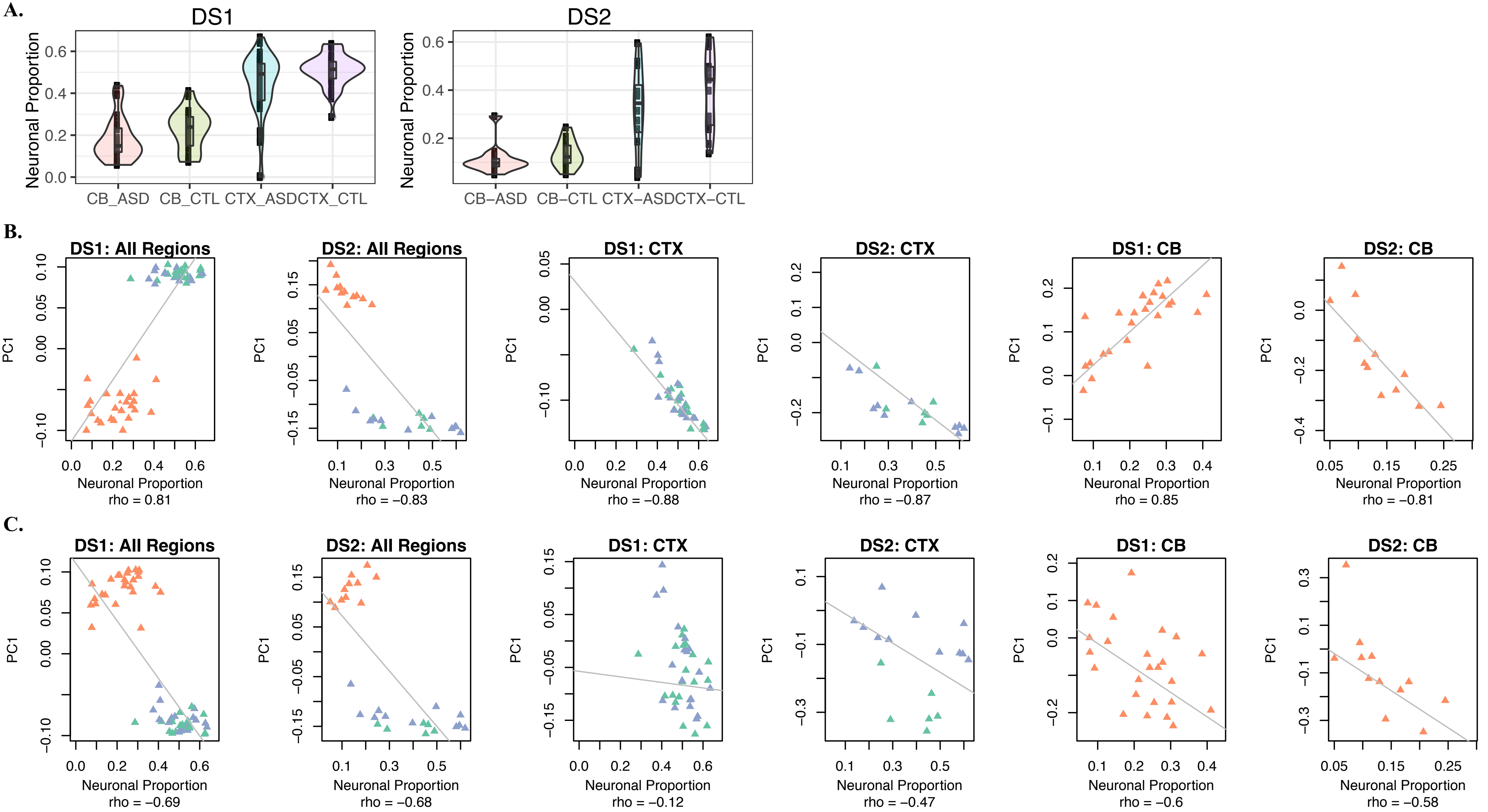


**Supplementary Figure 7. Estimated proportion of neurons in DS1 and DS2. (A)** Boxplots displaying the estimated proportion of neurons across brain regions and pheno-types, in DS1 (*left*) and DS2 (*right*). Boxplots were generated using the geom_boxplot and geom_violin functions in R; the horizontal line represents the median, boxes extend between the first and third quartiles, and whiskers extend to 1.5 IQR (inter-quartile range) from the box. Notches mark +/-1.58 IQR/sqrt(n), where n represents the number of data points. **(B)** Scatterplots of first principal component values of gene expression data (PC1, y-axis) vs. estimated proportion of neurons. **(C)** Scatterplots of first principal component values of circRNA expression data (PC1, y-axis) vs. estimated proportion of neurons. All neuronal proportion estimates are based on reference tran-scriptome data from Zhang et al. 2016.
