## Supplementary_Figure_8 for "The Landscape Of Circular RNA Expression In The Human Brain"

**A.**

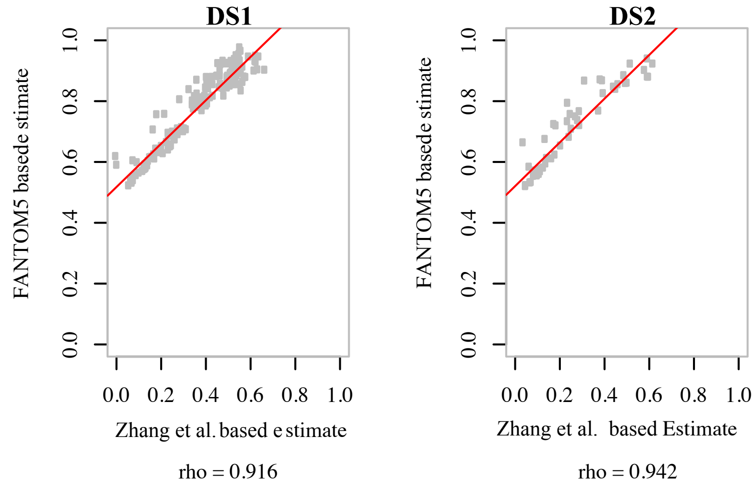


**B.**

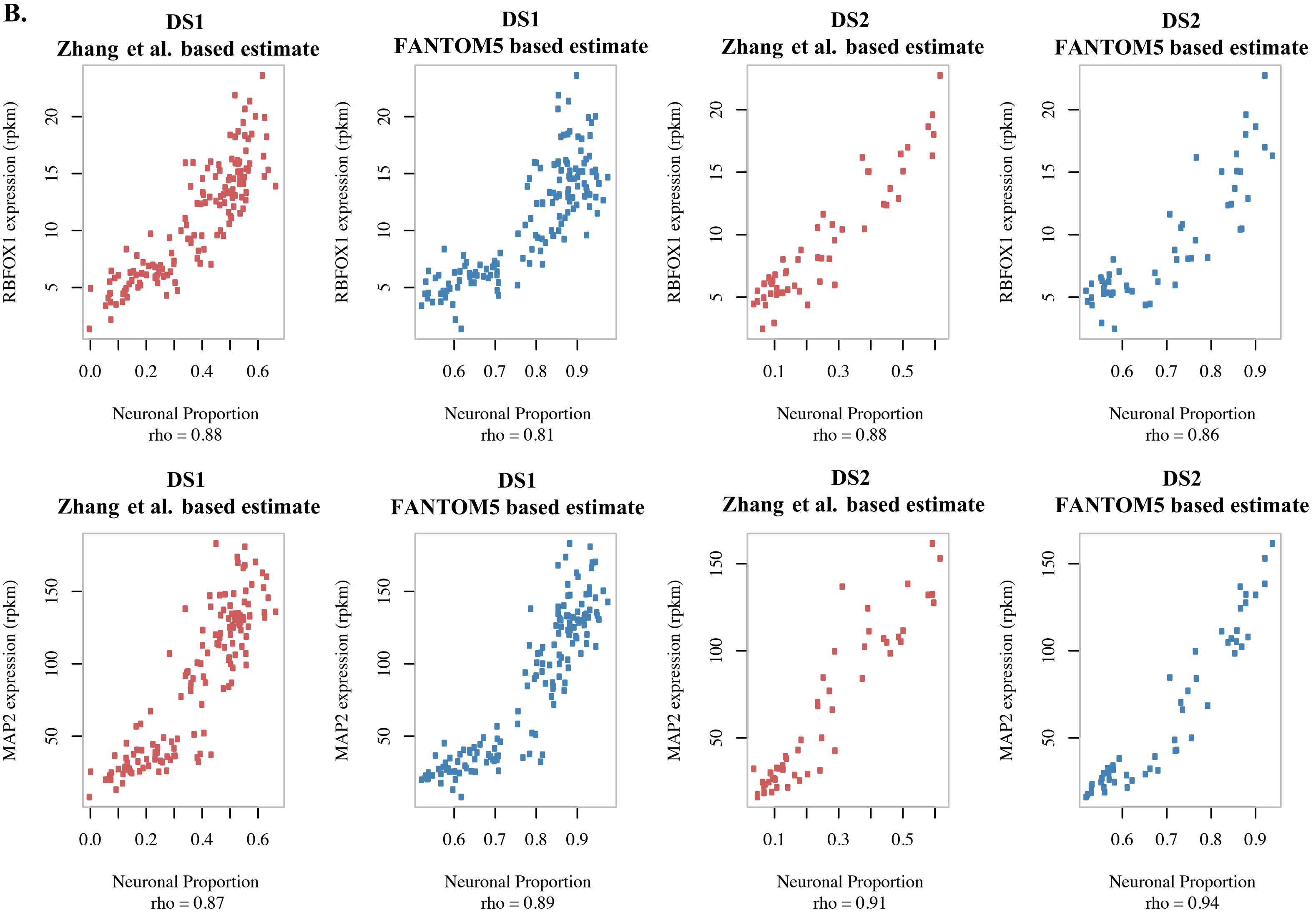


**Supplementary Figure 8. Assessment of cellular composition estimates. (A)** Scatterplot of estimated neuronal proportions based on the FANTOM5 and Zhang et al. reference transcriptome data in DS1 and DS2; *rho:* Spearman correlation coefficient. **(B)** Scatterplot of gene expression levels of two neuronal-specific genes (*RBFOX1*-top row, and *MAP2*-bottom row; y-axis) vs. estimated neuronal proportions (x-axis). *rho:* Spearman correlation coefficient.
